# Broad-spectrum RNA antiviral inspired by ISG15^-/-^ deficiency

**DOI:** 10.1101/2024.06.24.600468

**Authors:** Yemsratch T. Akalu, Roosheel S. Patel, Justin Taft, Rodrigo Canas-Arranz, Ashley Richardson, Sofija Buta, Marta Martin-Fernandez, Christos Sazeides, Rebecca L. Pearl, Gayatri Mainkar, Andrew P. Kurland, Rachel Geltman, Haylen Rosberger, Diana D. Kang, Ann Anu Kurian, Keerat Kaur, Jennie Altman, Yizhou Dong, Jeffrey R. Johnson, Lior Zhangi, Jean K. Lim, Randy A. Albrecht, Adolfo García-Sastre, Brad R. Rosenberg, Dusan Bogunovic

## Abstract

Type I interferons (IFN-I) are cytokines with potent antiviral and inflammatory capacities. IFN-I signaling drives the expression of hundreds of IFN-I stimulated genes (ISGs), whose aggregate function results in the control of viral infection. A few of these ISGs are tasked with negatively regulating the IFN-I response to prevent overt inflammation. ISG15 is a negative regulator whose absence leads to persistent, low-grade elevation of ISG expression and concurrent, self-resolving mild autoinflammation. The limited breadth and low-grade persistence of ISGs expressed in ISG15 deficiency are sufficient to confer broad-spectrum antiviral resistance. Inspired by ISG15 deficiency, we have identified a nominal collection of 10 ISGs that recapitulate the broad antiviral potential of the IFN-I system. The expression of the 10 ISG collection in an IFN-I non-responsive cell line increased cellular resistance to Zika, Vesicular Stomatitis, Influenza A (IAV), and SARS-CoV-2 viruses. A deliverable prophylactic formulation of this syndicate of 10 ISGs significantly inhibited IAV PR8 replication *in vivo* in mice and protected hamsters against a lethal SARS-CoV-2 challenge, suggesting its potential as a broad-spectrum antiviral against many current and future emerging viral pathogens.

**One-Sentence Summary:** Human inborn error of immunity-guided discovery and development of a broad-spectrum RNA antiviral therapy

## Introduction

Viral pathogens are a significant contributor to worldwide morbidity and mortality. The 1918 “Spanish Flu” is estimated to have killed 20-50 million Europeans (Bootsma and Ferguson, 2007; Erkoreka, 2010; Taubenberger and Morens). A hundred years later, despite vaccinations and available therapeutics, it is estimated that influenza virus continues to hospitalize about half a million individuals annually (Flerlage et al., 2021). Most recently, the SARS-COV-2 pandemic has resulted in 10-15 million deaths (Adam, 2022). Given climate change and rapid globalization practices, another viral pandemic is imminent (Baker et al., 2022; Gössling et al., 2021; Thoradeniya and Jayasinghe, 2021). Therefore, we must develop effective broad-spectrum antivirals to protect us from known, emerging, and unknown viral pathogens (Ianevski et al., 2019; Oksenych and Kainov, 2022). Soon after their initial discovery as an antiviral factor, type I interferons (IFN-Is) were thought to be an ideal broad-spectrum therapeutic agent (Antonelli et al., 2015; Dianzani; Postic and Arroyo, 1984). Unlike type III interferons (IFN-III), a related cytokine that uses a similar signaling cascade and is showing promise (Jagannathan et al., 2021), clinical experience has demonstrated limited utility for IFN-I in the treatment and prevention of acute viral infections (Sleijfer et al., 2005). As a potent inducer of inflammation, IFN-I is poorly tolerated with significant side effects (Aricò et al., 2019; Loftis and Hauser, 2004; Sleijfer et al., 2005). Recently, the therapeutic efficacy of IFN-I was shown to be disrupted by its induction of various cellular death pathways following the therapy (Karki et al., 2022). However, modifying the quantity and delivery methods might improve this (Monk et al., 2021). In addition, IFN-I is self-limiting therapeutically because, among the over 500 interferon-stimulated genes (ISGs) that act in concert to restrict viral replication, there are several negative regulators (Arimoto et al., 2018; Ivashkiv and Donlin, 2014). While essential in the context of natural infection to limit inflammatory damage to the host, these regulators are a hurdle to IFN-I therapeutic efficacy (Ivashkiv and Donlin, 2014). Their function creates a refractory state that prevents ISG expression in response to sustained stimulation (Makowska et al., 2011; Sugawara et al., 2021). In the shutdown of IFN-I activity, ubiquitin-specific peptidase 18 (USP18), in complex with IFN-I-stimulated gene 15 (ISG15), plays a prepotent role (Bogunovic et al., 2013; Hermann and Bogunovic, 2017; Honke et al., 2016; Perng and Lenschow, 2018). Upon expression, USP18 suppresses IFN-I signaling by physically displacing Janus Kinase 1 (JAK1) from the IFN-α receptor (IFNAR) complex (Malakhova et al., 2006), while ISG15 functions to stabilize USP18, protecting it from proteasome-mediated degradation (Vuillier et al., 2019; Zhang et al., 2015). As a result, deficiency for either ISG15 or USP18 cripples the system’s negative regulatory capacity, leading to persistent expression of ISGs and consequences for human health (Taft and Bogunovic, 2018). ISG15 deficiency results in partial loss of USP18 activity, leading to a mild form of persistent ISG expression. This has been observed in individuals from over 10 ISG15^-/-^ families, many of whom are well into adulthood (Bogunovic et al., 2012; Martin-Fernandez et al., 2020; Taft and Bogunovic, 2018; Zhang et al., 2015). On the other hand, a complete loss of USP18 results in a much starker clinical phenotype with a severe perinatal onset (Alsohime, 2020; Martin-Fernandez et al., 2022; Taft and Bogunovic, 2018). These two deficiencies thus point to safe and detrimental levels of IFN-I mediated inflammation (Jiménez Fernández et al., 2020; Taft and Bogunovic, 2018). Underscoring the protective potential of safe levels of IFN-I mediated inflammation, our group previously demonstrated that persistent ISG expression in immortalized ISG15^-/-^ patient-derived dermal fibroblasts resulted in greater control of a broad spectrum of human pathogenic RNA and DNA viruses (Speer, 2016). In these *in vitro* experiments, we mimicked the human ISG15-deficient ex vivo phenotype by first stimulating cells with IFN-I and then allowing them to rest before viral infection (Speer, 2016). At this point, in control cells, ISG15 and USP18 have shut down most ISG transcription. However, cells derived from ISG15^-/-^ patients with incomplete negative feedback retain expression of a subset of ISGs at approximately 1% of maximum expression. This is akin to the levels of ISGs in the blood of ISG15-deficient individuals (Speer, 2016). Cells were then virally challenged at a range of multiplicities of infection (MOIs) with a panel of RNA and DNA viruses (Speer, 2016). Across all viral infections tested, ISG15^-/-^ patient fibroblasts restricted infection significantly more than healthy control fibroblasts, sometimes by several orders of magnitude (Speer, 2016).

## Results

### ISG15 deficiency inspires the identification of a nominal collection of 10 ISGs with broad-spectrum antiviral potential

Inspired by the remarkable ability of ISG15-deficient cells to restrict a broad spectrum of viral infections with only a small fraction and low intensity of the ISG arsenal, we sought to define, formulate, and develop a therapeutic combination of ISGs capable of similar antiviral activity. ISG15^-/-^ individuals live with constant, low-level ISG expression in their blood (Zhang et al., 2015). This is due to tonic, environmental, and commensal-induced IFN-I, which they incompletely control. In sterile culture conditions, ISG15^-/-^ cells do not experience baseline ISG expression. To recapitulate the *in vivo* effect of defective negative feedback *in vitro*, we stimulated cells with IFN-I for 12 hours, then washed off the IFN-I and incubated the cells for another 36 hours. By this point, 48 hours after IFN-I exposure, USP18, stabilized by ISG15, has downregulated most ISG transcription in fibroblasts from healthy controls—a process that is incomplete in ISG15^-/-^ individuals (Figure 1A). As IFN-I primarily induces ISG15 and USP18, their absence does not impact initial IFN-I activation or peak ISG expression levels. However, ISG15-deficiency results in persistent, low-level ISG expression, as demonstrated by *IFIT1*, *BST2*, *RSAD2*, *MX1* and *SIGLEC1* mRNA levels (Figure 1A). Remarkably, this residual ISG expression, present at only ∼1 % of peak IFN-I induced activity, is sufficient to substantially restrict a broad-spectrum of viral agents (Speer, 2016). We have now assessed the impact of persistent ISG expression in ISG15^-/-^ human immortalized fibroblasts using a Zika virus (ZIKV) infection model (Figure 1B). An overnight ZIKV challenge at an MOI of 1 in IFN-I exposed fibroblasts was inhibited 14-fold in ISG15^-/-^ compared to similarly treated healthy control cells (Figure 1B, left), and this inhibition translated to less productive virus formation (Figure 1B, right). To precisely identify the ISGs responsible for broad-spectrum antiviral activity, we performed RNA sequencing on fibroblasts from ISG15^-/-^ and healthy individuals under two conditions: unstimulated and IFN-I exposed (12-hour IFN-I stimulation followed by 24-hour rest). ISG15^-/-^ fibroblasts harbor either a nonsense or frame-shift mutation at position c.379 or c.336_337, respectively. While these mutations result in total loss of protein expression, their location in the last exon of *ISG15* circumvents RNA decay (Bogunovic et al., 2012; Zhang et al., 2015), allowing detection of the transcript by RNA sequencing. Difference-of-differences gene expression analyses identified 61 ISGs that were differentially responsive to IFN-I between ISG15^-/-^ and control cells (Figure 1C). Additionally, we noticed that a small subset of the differentially regulated ISGs in ISG15^-/-^ cells also remained elevated above baseline in the prime-rested fibroblasts from healthy controls, albeit at lower levels (Figure 1D). Notably, IFN-I exposed ‘wild-type’ controls retain some antiviral protection (Figure 1B), suggesting that this antiviral function could be ISG expression dose-dependent. Thus, we hypothesized that the ISGs with lingering expression in a ‘wild-type’ background evolved lower shutdown kinetics because maintaining low-grade expression of a subset of ISGs is sufficient to exert some antiviral effects (Figure 1B). Therefore, heightened expression levels of select ISGs may, at least partially, underlie the improved resistance in IFN-I exposed ISG15^-/-^ cells (Figure 1D). Based on these observations and the antiviral function that each ISG is known to have at distinct steps of the viral life cycle, we chose 10 persistently elevated ISG effectors (Figure 1D-F). We posited that they were sufficient to replicate broad-spectrum viral resistance documented in ISG15-deficiency. These 10 ISGs could confer broad-spectrum antiviral resistance similarly to HIV treatment with HAART (Highly Active Antiretroviral Therapy), which targets multiple stages of the viral lifecycle (Figure 1E-F).

**Figure 1.**
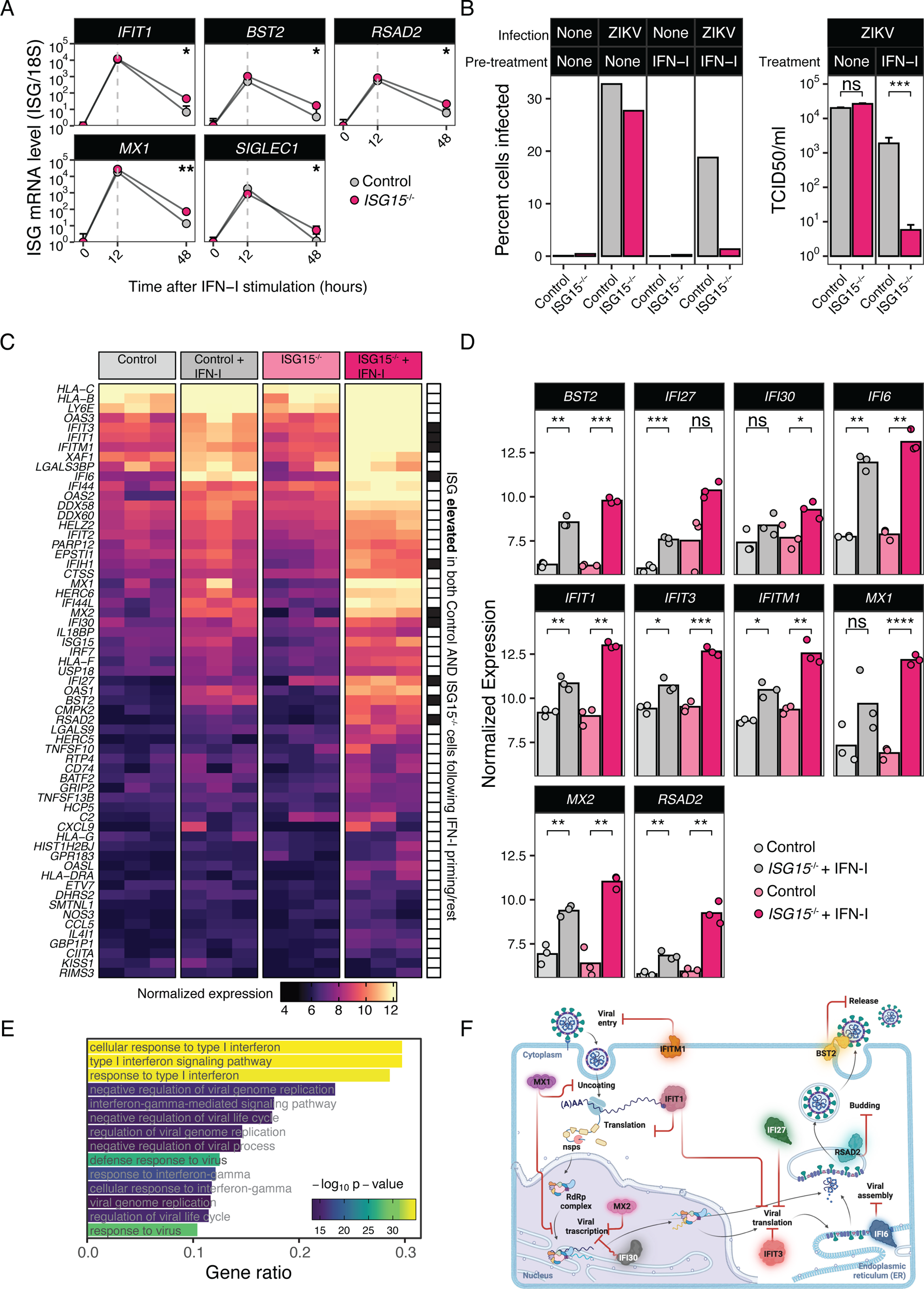
ISG15 deficiency inspires the identification of a nominal collection of 10 ISGs with broad-spectrum antiviral potential. **(A)** Relative mRNA level time-course quantification of *IFIT1, BST2, RSAD2, MX1,* and *SIGLEC1* from ISG15^-/-^ patients (n=3) or control (n=3) hTERT-immortalized fibroblasts treated with 1,000 IU ml^−1^ IFN-α2b for 12 h, washed and allowed to rest for 36-hour post-treatment. *p<0.05, **p<0.01, one-sided paired Student’s t-test**. (B;** left) At the 48-hour time point, cells were infected with Zika virus (PRVABC59) at a multiplicity of infection (MOI) of 1.0 overnight, fixed, subjected to a live/dead and viral staining for Zika virus envelope protein (4G2 Antibody), and processed and analyzed by flow cytometry. The percentages of infected cells was quantified. **(B;** right) Viral titers were from supernatants from Zika virus-infected cells were quantified by TCID_50_ on control fibroblasts. n.s. (not significant), ***p<0.001 by one-sided paired Student’s t-test**. (C)** Differentially IFN-I responsive genes between unstimulated and 12-hour IFN-I stimulated, 24-hour rested fibroblasts from control and ISG15^-/-^ individuals. Heatmap color intensity depicts variance stabilized, normalized expression values. Row annotation bar describes overlap of elevated ISGs in both ISG15^-/-^ and control IFN-I exposed conditions (relative to unstimulated; genotype matched). **(D)** Normalized expression values of 10, select, ISGs elevated in both ISG15^-/-^ and control IFN-I exposed conditions, compared to respective unstimulated controls. *IFI30* and *MX1* did not meet significance cut-off but were included due to their antiviral functions. *IFIH1* is excluded due to its role as a dsRNA sensor and signal potentiator of the IFN-I response. *p_adj_<0.05, **p_adj_<0.01, ****p_adj_<0.0001 by Benjamin-Hochberg method. **(E)** Gene ontology (GO) enrichment analysis of differentially IFN-I responsive genes indicated in (C). Gene ratio indicates the proportion of genes in the input list that are associated with each GO term compared to the total number of genes shown in (C). **(F)** Literature-defined schematic and hypothetical combinatorial antiviral activity of 10-ISG collection.

### Select persistently elevated ISGs expressed in isolation show known but limited antiviral restriction

To test the antiviral potential of these 10 ISGs, we stably transduced each ISG into a JAK1^-/-^, IFN-I non-responsive fibrosarcoma (U4C) cell line (Figure 2A). Leveraging the inability of these cells to respond to IFN-I enabled us to assess the antiviral capacity of each ISG in question, independent of any endogenous IFN-I produced due to viral challenge (Figure 2A). After validating ISG expression in their respective cell lines (Figure 2B), we tested the antiviral potency of these ISGs against a panel of RNA viruses: Vesicular Stomatitis Virus (VSV), Influenza A virus (IAV PR8), and Zika Virus (ZIKV PRVABC59). As expected, MX1 was sufficient to restrict VSV and IAV PR8 infection (Figure 2C-D), while IFI6 proved capable against Zika Virus (Figure 2E). While IFI6 showed antiviral activity against ZIKV, infection was controlled to even greater levels by IFN-I exposed fibroblasts from ISG15^-/-^ individuals (Figure 1B). Observing that IFN-I ISG15^-/-^ fibroblasts outperformed the established pan-flavivirus restriction factor IFI6 in a ZIKV challenge led us to postulate that the combination of the 10 ISGs may work cooperatively to restrict viral infection.

**Figure 2.**
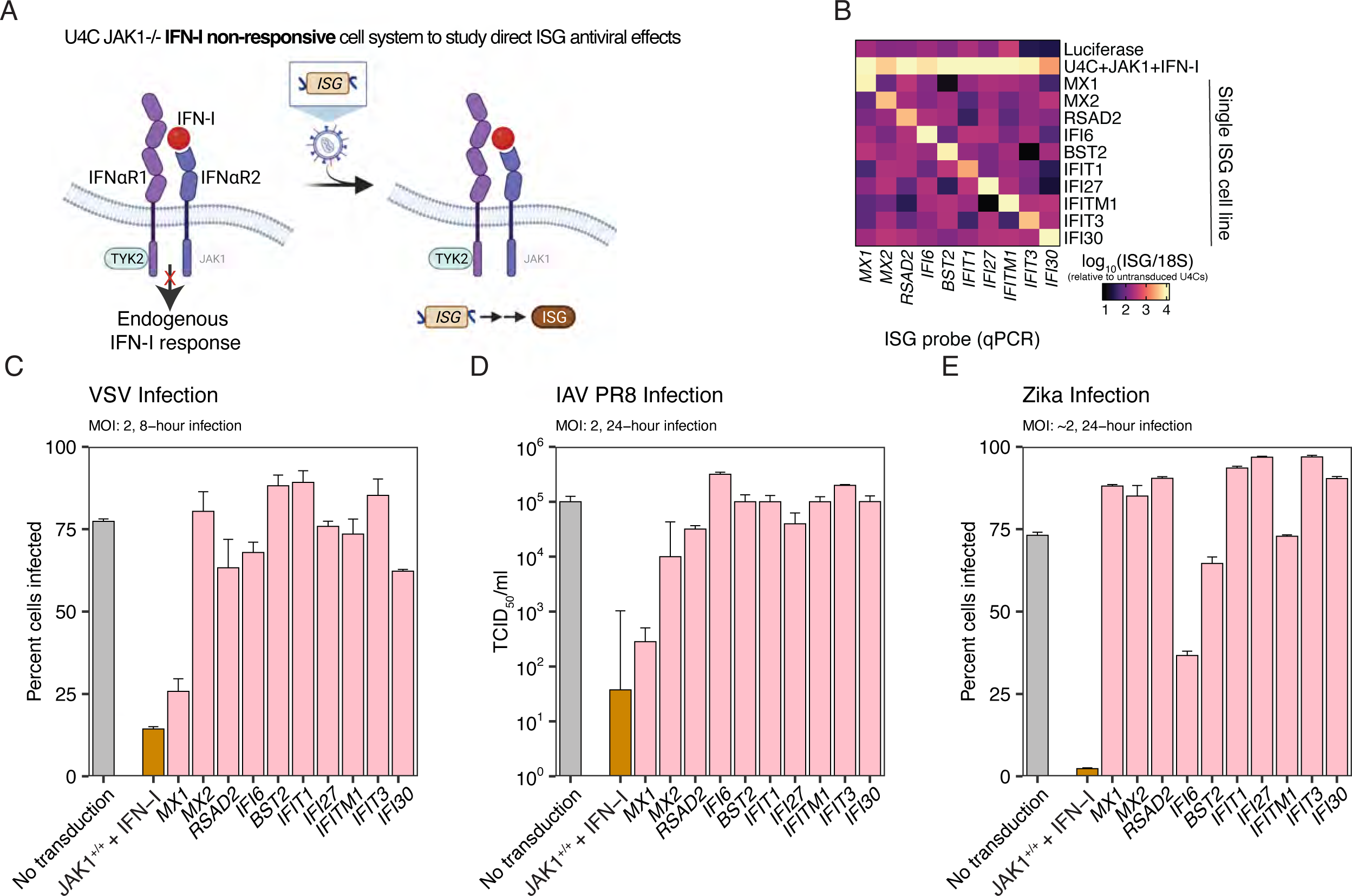
Select persistently elevated ISGs expressed in isolation show known but limited antiviral restriction. **(A)** Schematic of *in vitro* JAK1^-/-^ IFN-I non-responsive cell system to study direct ISG antiviral effects. Cells are transduced with lentiviruses, encoding an individual ISG (from 10-ISG collection) or luciferase or red fluorescent protein (RFP) prior to viral challenge. **(B)** Relative qPCR quantification of mRNA transcripts of encoded ISGs in untransduced and lentivirus-transduced lines. **(C)** Lentivirus transduced cells were subject to viral challenge with vesicular stomatitis virus (MOI: 2; 8-hour infection), **(D)** influenza A virus (MOI: 2; 24-hour infection), and **(E)** Zika virus (MOI: ∼2, 24-hour infection). Bars represent average infection values between 3 biological replicates.

### Synthetically modified RNA delivery of 10-ISG collection is sufficient to restrict viral infection in vitro

To interrogate this, we adopted a synthetic, modified RNA (modRNA) technology coupled with lipid nanoparticle (LNP) delivery to introduce a collection of 10 ISGs (Table S1) into cells before viral challenge (Figure 3A). Using a modRNA-LNP encoding emerald GFP, we detected high transfection efficiency in U4C *JAK1^-/-^*IFN-I non-responsive cells and confirmed that modRNA encoding GFP is translatable (Figure 3B). To assess the efficiency and immunogenicity of modRNA-LNP: 10-ISG transfection, we measured modRNA levels of our target ISGs 16 hours post-transfection (Figure 3C). Quantifying levels of target ISG expression, we detect mRNA of all 10 ISGs (Figure 3C), and the level of ectopic modRNA ISGs detected is comparable to peak expression induced by IFN-I stimulation (Figure 3C). Expression of GFP and all 10 ISGs at the protein level was also confirmed using either mass spectrometry or western blotting (Figure S1A-B). Additionally, to demonstrate the specificity of modRNA-LNP: 10 ISG delivery, we confirmed that the expression of USP18, an ISG not included in the 10-ISG combination, remained unchanged compared to the untreated and luciferase-transfected conditions. Similarly, delivery of modRNA-LNP encoding mouse ISG orthologues into human-derived HEK293T cells did not result in the induction of human *USP18*, *ISG15* and *IFI44L*—ISGs unrelated to the delivered subset which were potently induced by IFN-I stimulation (Figure S2A). Consistently, induction of IFN-I, IFN-II or IFN-III was not observed in U4C, A549 and 2fTGH cells transfected with the 10-ISGs (Figure S2D-F). In parallel, we tested if delivery of the 10ISGs drives a refractory state that halts subsequent activation of the IFN-I system. A549 cells were mock-treated, transfected with 10ISGs or stimulated with IFN-I and rested before a second stimulation with IFN-I. As expected, we found that cells pre-treated with IFN-I entered a refractory state, evidenced by the accumulation of USP18 and downregulation of p-STAT1 and pSTAT2 (Figure S3). In contrast, cells primed with modISG10 responded to the second IFN-I stimulation with a dose-dependent induction of pSTAT1 and pSTAT2, resembling mock-treated cells in their USP18 levels. Taken together, these observations suggested that modRNA was introducing the effector function of ISGs without the side effects of IFN-I. Next, to assess whether modRNA-LNP delivery of ISGs was sufficient to restrict viral infection, we took cells transfected with single ISGs or the 10-ISG collection and challenged them with VSV (Figure 3D). As expected, and previously demonstrated, cells transfected with *MX1* modRNA exhibit VSV infection restriction levels similar to, if not greater than, those of JAK1-complemented cells pre-stimulated with IFN-I. No other single ISG transfections appeared to control infection relative to the non-transfected control. However, transfection with the 10-ISG collection controlled infection to similar levels as cells pre-stimulated with IFN-I, (Figure 3D). To assess the combinatorial antiviral potential of the 10-ISG collection, we revisited our ZIKV infection model. Considering our observation of a moderate restriction of ZIKV by IFI6 (Figure 2E) and the modular nature of the modRNA-LNP cocktail generation process, we added an additional transfection condition to deliver 9 ISGs (excluding *IFI6*). As expected, the 10-ISG combination and IFI6 alone restricted ZIKV infection (Figure 3E). Interestingly, the 9 ISG combination cocktail, in the absence of IFI6, also appreciably restricted ZIKV infection, suggesting the other ISG effectors act in concert by unknown mechanisms to restrict infection (Figure 3E). Additionally, we tested the efficacy of the 10 ISG collection against another flavivirus, West-Nile virus (WNV). We found that pretreatment with modISG10, but not modIFI6 or modGFP, significantly protected 2fTGH cells from WNV infection (Figure S4A). Similarly, pre-treatment with modISG10 protected B6 MEFs from WNV infection, similar to treatment with IFN-I (Figure S4B). While treatment with modIFI6 noticeably reduced the percentage of infected cells compared to mock controls, its effect was similar to that of modGFP, suggesting that any benefit was a result of nonspecific inflammation trigged by the delivery of modRNA in these cells (Fig S4B). Next, we further explored the antiviral capacity of our 10-ISG collection to restrict SARS-CoV-2 infection (Figure 3F). Single ISG modRNA-LNP transfections had minimal to no effect on restricting viral infection of IFN-I non-responsive JAK1-/-fibrosarcoma cells. A dramatic antiviral effect was observed only when the 10 ISGs were expressed in combination (Figure 3F). To test if the antiviral effect observed following delivery of the 10 ISGs is accompanied by cell death, we transfected fibroblasts with LNP alone, modGFP, modIFI6, modISG10 or treated them with IFN-I. There were no detectable differences in cell viability among all conditions at 24 and 48hours following treatment (Figure S5A-B). The total percentage of live cells or cleaved caspase 3-positive cells across the tested conditions was comparable to cells left in media alone (Figure S5A-B). Only fibroblasts incubated with cell death inducing agents, Shikonin and 10% DMSO, showed a marked decline in cell viability. Combined, these data suggest that the 10 modRNA combination restricts VSV, IAV, ZIKV, WNV and SARS-CoV2 infection in *vitro*.

**Figure 3.**
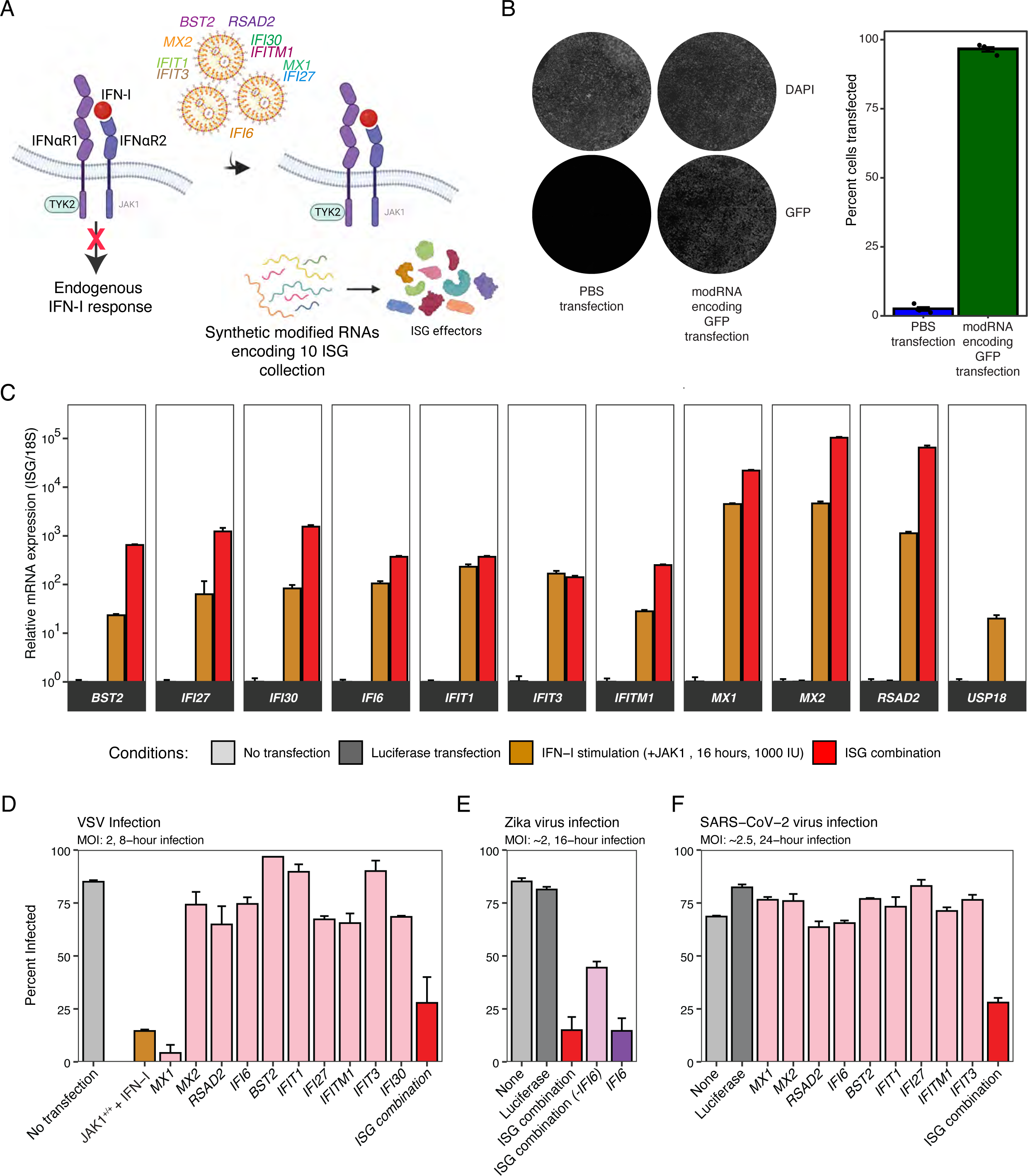
Synthetically modified RNA delivery of 10-ISG collection is sufficient to restrict viral infection. **(A)** Schematic of synthetic modified RNA (modRNA) delivery in IFN-I JAK1^-/-^ non-responsive cell system. Modified RNAs encoding emerald GFP or the 10-ISG collection are pooled in an equimolar ratio, packaged into lipid nanoparticles (LNPs), and transfected into cells for 16 hours before imaging, relative mRNA quantification, and viral challenge. **(B)** Representative brightfield and GFP images of cells treated with PBS or modRNA-LNP encoding emerald GFP, 16 hours post-transfection (left). Quantification of GFP^+^ cells (right). **(C)** Relative qPCR quantification of exogenously expressed 10 ISGs and *USP18* in non-transfected, luciferase modRNA-LNP transfected, JAK1-complemented and IFN-I pre-stimulated, and 10-ISG collection modRNA-LNP transfected cells, 16 hours post-transfection. USP18 levels were measured to assess non-specific IFN activity of modRNA-LNP administration. **(D)** ModRNA-LNP transfected cells were subject to viral challenge with vesicular stomatitis virus (MOI: 2; 8-hour infection), **(E)** Zika virus (MOI: ∼2, 24-hour infection), and **(F)** SARS-CoV-2 virus (MOI: ∼2.5; 24-hour infection). For SARS-CoV-2 infections, U4C JAK1^-/-^ IFN-I non-responsive cells were transduced with human ACE2 to make cells susceptible to viral infection. Bars represent average infection values between 3 biological replicates.

### Synthetically modified RNA intranasal delivery of 10 ISG collection is sufficient to restrict IAV and SARS-CoV-2 in vivo

Having validated the potency of the 10-ISG collection against multiple viruses *in vitro*, we set out to further assess the ability of these ISGs to restrict infection *in vivo*. We first tested the antiviral efficacy of the 10-ISG collection in an *in vivo* mouse model of IAV PR8. Mice were treated with PBS, modRNA-LNP encoding mouse ISG orthologues (Table S3, Figure S6) or emerald GFP, on day -1 and challenged with a lethal dose (5× LD50) of IAV PR8 on day 0 (Figure S7A). The mice were monitored for viral titers at D3, loss of body weight and survival. Mice that received either modMx1 or modISG10 had significantly lower viral burden, estimated by measuring viral titer and plaque size, in their lungs at D3 post-infection, compared to mice in the modGFP group (Figure S7D-E). Administration of modMx1 or modISG10 controlled lung viral load better than IFN-I. While prophylactic treatment of mice with the 10-ISG collection was not able to rescue survival at a high dose of viral challenge, it was sufficient to significantly reduce IAV PR8 replication in the lung. Next, we adopted the Syrian Golden Hamster (*Mesocricetus auratus*) model to evaluate SARS-CoV-2 infection (Figure 4A). We first assessed the efficiency of two different administration routes for delivering modRNA-LNPs to animal lungs: intranasal and intratracheal (Figure 4A). Harvesting lung tissue from animals, we observed appreciable efficiency of emerald GFP expression in the lungs of animals in both the intranasal and intratracheal delivery groups (Figure 4B-C). Next, to assess the antiviral capacity of the 10-ISG combination delivered intranasally, we treated animals with PBS, modRNA-LNP encoding hamster ISG orthologues (Table S2, Figure S6) or emerald GFP, 1 day before SARS-CoV-2 viral challenge. Animal weights were monitored, and select tissues were collected at days 2 and 4 post-infection (Figure 4A). Initially, all animals gained weight; however, beginning at day 2 post-infection, experimental groups began to stratify (Figure 4D). By day 7 post-infection, compared to PBS-treated animals or animals treated with modRNA-LNP encoding emerald GFP, animals treated with modRNA-LNP encoding the 10 ISGs were protected from sickness associated weight loss (Figure 4D). Furthermore, viral titer measurements in the lungs at day 2 post-infection revealed a reduction in the levels of infectious SARS-CoV-2 virions in the lungs of the 10-ISG combination treatment group compared to mock and emerald GFP groups (Figure 4E, left). Similar differences in viral burden were not detected in the olfactory bulb at the same time point, likely due to the restricted delivery of modISGs to the lungs. Furthermore, in transcriptomic analyses of day 4 SARS-CoV-2 infected hamster lungs, we detect expression of viral transcripts in mock-treated and emerald GFP expressing control animals. In contrast, detection of viral transcripts is relatively lower in the animals treated with the 10-ISG collection (Figure 4F). This is accompanied by a stark difference in host gene expression profiles between animals treated with 10-ISG collection and control groups (Figure 4F-G). Principle component analysis of the infected hamster lungs clustered PBS- and GFP modRNA treated samples closely together and far away from the ISG10 modRNA samples (Figure 4G). Clinical scoring of infected hamsters’ lungs, following hematoxylin and eosin staining on day 7 post-infection, revealed extensive disease pathology, notably indicated by airway constriction (Figure 4H). Overall, animals assigned to modISG10 group had lower pathology scores than control hamsters, suggesting that modRNA-LNP administration of the 10-ISG collection is sufficient to restrict SARS-CoV-2 infection and prevent lung inflammation and damage.

**Figure 4.**
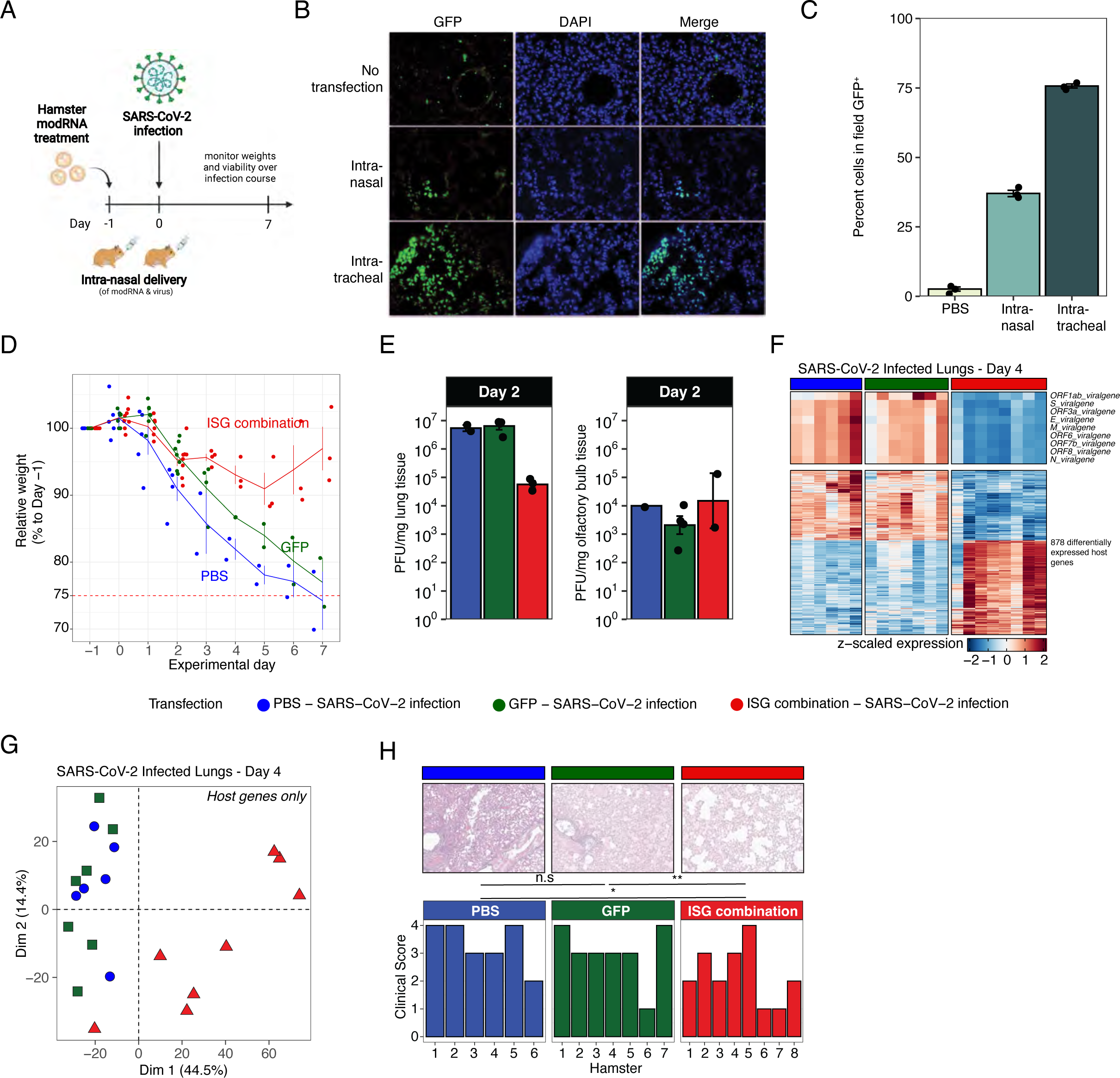
Synthetically modified RNA intranasal delivery of 10 ISG collection is sufficient to restrict SARS-CoV-2 *in vivo*. **(A)** Experimental design of intranasal modRNA-LNP delivery and SARS-CoV-2 viral challenge in the Syrian golden hamster model. **(B)** Representative confocal microscopy images of day 2 lung sections from modRNA-LNP: emerald GFP-treated or PBS-treated hamsters. **(C)** Quantification of (B) showing GFP^+^ cells per field (n=3 animals/group). **(D)** Weight loss curves over 7-day infection time course of PBS-treated, modRNA-LNP: emerald GFP-treated, and modRNA-LNP: 10-ISG collection-treated hamsters following SARS-CoV-2 infection. The dotted red line represents the weight loss threshold per the predefined humane endpoint (n=3 animals/group). **(E)** Day 2 SARS-CoV-2 viral loads detected in PBS-treated or modRNA-treated hamster lungs (left) and olfactory bulbs (right) (n=3 animals/group). **(F)** Viral gene expression and differentially expressed host genes detected between PBS-treated or modRNA-treated hamster lungs on day 4 post-infection (n=6-8 animals/group). **(G)** Principal component analysis on host gene expression striates modRNA-LNP: 10-ISG collection treated hamsters from PBS-treated and modRNA-LNP: emerald GFP hamsters’ groups (n=6-8 animals/group). **(H)** Clinical, histological scoring of PBS or modRNA-LNP treated hamster lungs at day 7 post-infection (n=6-8 animals/group).

## Discussion

None of the 67 antiviral drugs currently approved by the FDA are clinically used as an effective broad-spectrum treatment for viral infections (De Clercq and Li, 2016). Some viral infections are self-resolving, but many cause severe morbidity and mortality. In 2014, the Ebola virus killed 11,000 people in West Africa (2020). Since 2015, infections by ZIKV in the Americas has resulted in thousands of infants being born with microcephaly (2014). Just under 100 years ago, the Spanish Flu was estimated to have killed 20-50 million Europeans (Taubenberger and Morens). Even today, flu kills 3,000-50,000 individuals annually in the US alone, depending on the flu strain, with the elderly and children most affected (Baguelin et al., 2013; Sullivan et al., 2019; Taubenberger and Morens). Recently, we have witnessed an unprecedented global pandemic caused by SARS-CoV-2, which has claimed millions of lives within the last few years (Kissler et al., 2020). History shows that every two to three years, there is an epidemic of a highly contagious, often deadly, viral disease with tremendous associated morbidities (Piret and Boivin, 2021; Tognotti, 2013). As a society, we are not prepared for future outbreaks of either novel strains of existing viruses (*e.g.,* SARS-CoV-2 or *a* repeat of the 1918 Spanish-like flu) or an unknown highly pathogenic virus. The availability of a broad-spectrum antiviral drug would significantly increase our ability to fight off known and unknown viral diseases alike. By developing the concept of a broad-spectrum antiviral drug effective against many existing viruses, we are expanding our arsenal against unknown, potentially highly pathogenic viruses. A drug targeting the flu could save 3,000-50,000 lives annually in the US, while a drug effective against the ZIKV could prevent significant morbidities. Additionally, a drug effective against SARS-CoV2 would have saved millions of lives. The development of such drugs would be a significant leap forward, well worth the effort in terms of our readiness to deal with unknown viruses or biological warfare. Genetic studies in humans with inborn errors in the IFN-I system have manifested how indispensable this system is for fighting viral infections (Casanova and Abel, 2021; Taft and Bogunovic, 2018). In contrast, examples of overt IFN-I signaling due to genetic defects in controlling production or response to IFN-I also define how detrimental prolonged, excessive IFN-I signaling can be (Casanova and Abel, 2021; Duncan et al., 2021; Sancho-Shimizu et al., 2011; Zhang et al., 2020). ISG15^-/-^ patients have likely defined the optimal “Goldilocks zone” of safe levels of IFN-I signaling levels: sufficient for enhanced control of every virus tested, with minimal side effects (isolated, self-resolving, early childhood skin lesion, and BCG vaccine lymphadenopathy). Notably, most of these individuals are well into adulthood (Taft and Bogunovic, 2018). Herein, inspired by these patients’ impressive ability to restrict a broad-spectrum of viral infections with only a small fraction of the ISG arsenal, we defined a nominal but sufficient syndicate of ISGs that can achieve similar antiviral activity. In line with the need for a broad-spectrum antiviral, we leveraged recent advances in RNA therapeutics and lipid nanoparticle technology to deliver synthetic modRNAs expressing the ISG combination *in vitro* and *in vivo*. RNA delivery of the ISG combination was not only lowly immunogenic, highly expressed, and well-tolerated *in vitro* and *in vivo* but also restricted infection of SARS-CoV-2 in hamsters. Ultimately, we engineered a system where we have further reduced the already small ISG repertoire to a 10 ISG effector syndicate that effectively controls viral infection and severity of viral disease. With this, we aimed at extracting the essence of antiviral effects while reducing the inflammation by omitting the hundreds of naturally induced ISGs. As such, the modality of transient gene therapy where inhalation of a syndicate of ISG modRNAs is poised to serve as a true broad-spectrum antiviral to be used against an existing or yet unknown respiratory virus, with a la carte reduction of perhaps 1 to 3 select ISGs for specific infections.

## Acknowledgments

We thank Thomas Moran and the Center for Therapeutic Antibody Discovery at the Icahn School of Medicine at Mount Sinai for kindly providing anti-SARS-CoV2 nucleocapsid antibody. We thank all members of the laboratory of Brad Rosenberg and of Dusan Bogunovic for advice, helpful discussions, and support. This work was partly supported by NIH grants R01 AI151029 and U01 AI150748 and funds from the Department of Microbiology, Icahn School of Medicine at Mount Sinai. We also acknowledge the Biorender scientific illustration suite which was used to generate many descriptive illustrations presented in this work.

## Funding

This research was supported by National Institute of Allergy, and Infectious Diseases (NIAID) Grants R01AI151029, R01AI127372, R41AI164999, R21AI134366, and R21AI129827, and funding from the March of Dimes, awarded to DB, by NIH grants R01AI151029 and U01AI150748, and funds from the Department of Microbiology, Icahn School of Medicine at Mount Sinai, awarded to BRR, and R01AI150837 awarded to JKL. This research was also partly supported by CRIPT (Center for Research in Influenza Pathogenesis and Transmission), an NIAID supported Center of Excellence for Influenza Research and Response (CEIRR, contract #75N93021C00014 to RAA and AG-S), and by NIAID Grant U19AI135972 and DARPA Grant HR0011-19-2-319 0020 to AG-S. RSP is supported by Virus-Host T32 training grant T32 AI07647. The content is solely the responsibility of the authors and does not necessarily represent the official views of the funders, including the National Institutes of Health.

## Author contributions

Conceptualization: YTA, RSP, JT, DB

Methodology: YTA, RSP, JT, SB, AR, RC, GM, RLP, AAK, KK, JA, YD, LZ, JL, RAA

Investigation: YTA, RSP, JT, RC, MMF, CS, AK, RG, HR, DK, JA, RLP, RAA, DB

Visualization: YTA, RSP

Funding acquisition: LZ, JL, JJ, RAA, BRR, AGS, DB

Project administration: RAA, AGS, DB

Supervision: RAA, BRR, AGS, DB

Writing – original draft: RSP, JT, DB

Writing – revised draft: YTA, DB

## Declaration of Interests

DB reports ownership in Lab11 Therapeutics. A patent covering the usage of 10 interferon stimulated gene collection as a broad-spectrum antiviral has been filed by the Ichan School of Medicine at Mount Sinai. The A.G.-S. Laboratory has received research support from Pfizer, Senhwa Biosciences, Kenall Manufacturing, Avimex, Johnson & Johnson, Dynavax, 7Hills Pharma, Pharmamar, ImmunityBio, Accurius, Nanocomposix, Hexamer, N-fold LLC, Model Medicines, Atea Pharma, Applied Biological Laboratories and Merck, outside of the reported work. A.G.-S. has consulting agreements for the following companies involving cash and/or stock: Vivaldi Biosciences, Contrafect, 7Hills Pharma, Avimex, Vaxalto, Pagoda, Accurius, Esperovax, Farmak, Applied Biological Laboratories, Pharmamar, Paratus, CureLab Oncology, CureLab Veterinary, Synairgen, and Pfizer, outside of the reported work. A.G.-S. has been an invited speaker in meeting events organized by Seqirus, Janssen, and AstraZeneca. A.G.-S. is the inventor of patents and patent applications on antivirals and vaccines for the treatment and prevention of virus infections and cancer, owned by the Icahn School of Medicine at Mount Sinai, New York, outside of the reported work.

---

**Figure S1.**
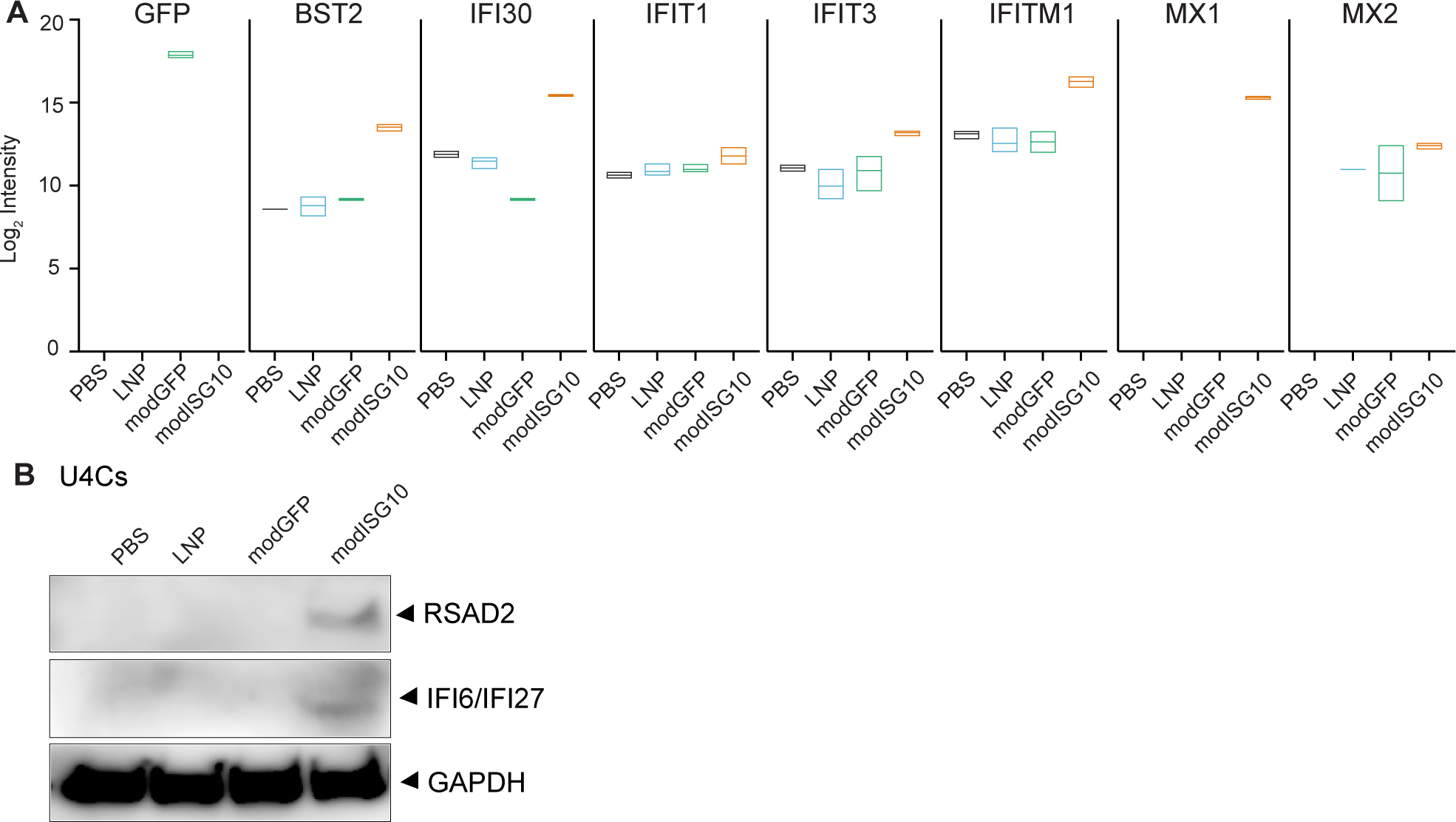
Efficient *in vitro* delivery of synthetically modified 10-ISG RNA detected at the protein level. (**A**) Levels of emerald GFP and seven different ISGs (BST2, IFI30, IFIT1, IFIT3, IFITM1, MX1 and MX2) were quantified in non-transfected U4Cs, or U4Cs transfected with RNAiMax Lipofectamine alone (LNP), modGFP RNA, or modISG10 RNA, as assessed by mass spectrometry. (**B**) RSAD2, IFI6/IFI27 proteins were detected by western blot of U4Cs transfected with modRNA of 10-ISG collection.

**Figure S2.**
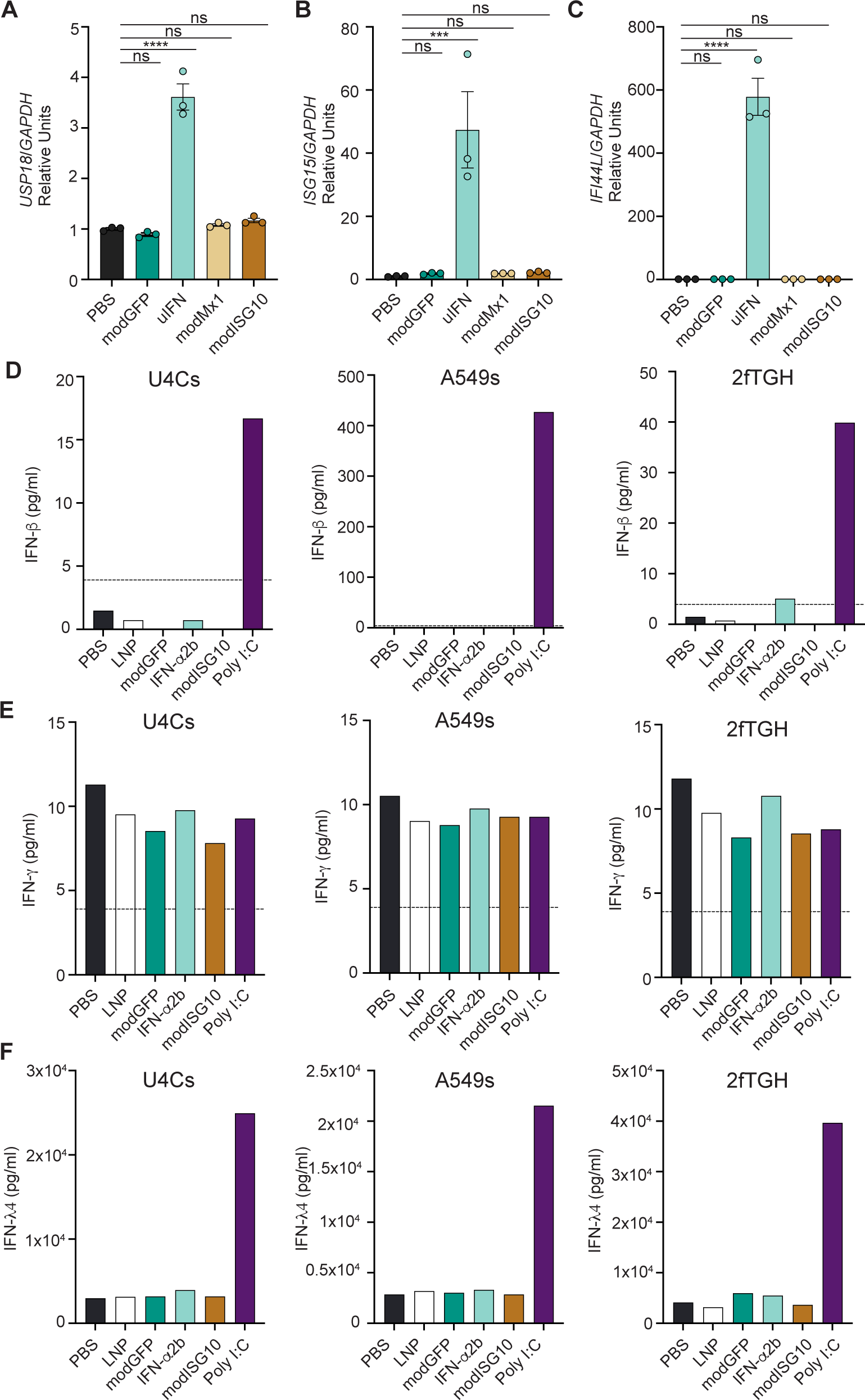
Delivery of packaged 10-ISG RNA *in vitro* does not trigger broader ISG or IFN-I/II/II production. (**A**) HEK293T cells were treated with PBS or transfected with modRNA-LNP encoding GFP, mouse *Mx1* or mouse ISG orthologues for the 10-ISG combination. Human *USP18*, *ISG15* and *IFI44L* mRNA were quantified by qPCR (n=3 samples/group). ***p<0.001, ****p<0.0001, one-way ANOVA Dunnett’s test. (**B**) U4Cs, A549s and 2fTGH cells were treated with PBS, RNAiMax Lipofectamine (LNP) alone, modGFP, IFN-α2b (100IU) or polyI:C (100μg/ml) for 24hrs. Levels of (**D**) IFN-β, (**E**) IFN-γ and (**F**) IFN-λ4 were then measured in the supernatants of these cells. Treatment with polyI:C served as a positive control for interferon production. Broken lines represent the detection limit for each cytokine indicated.

**Figure S3.**
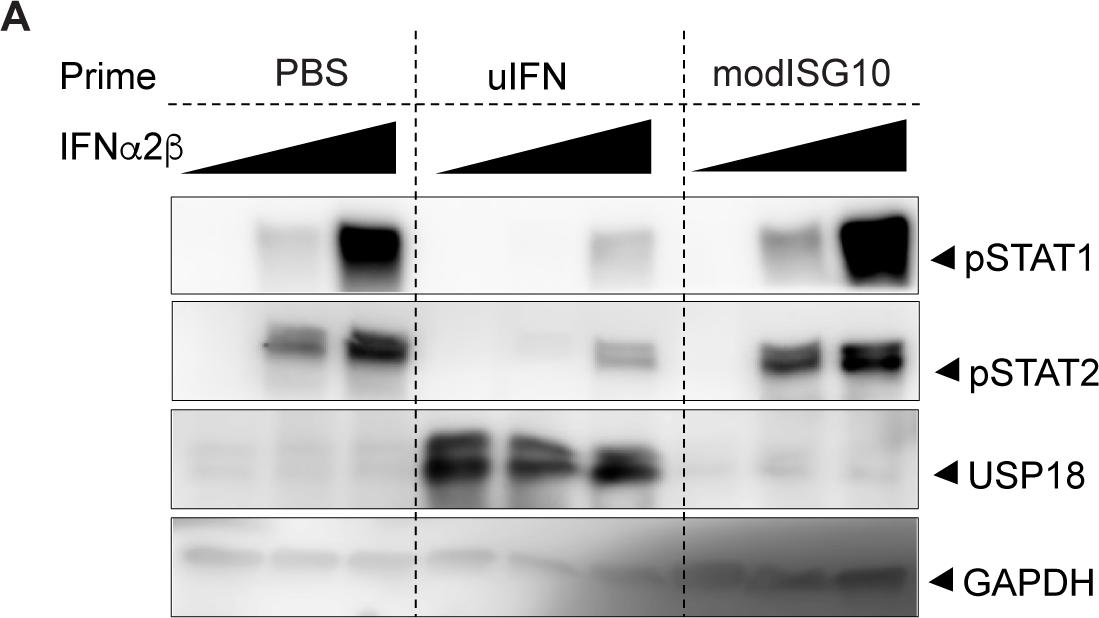
Following pre-treatment with 10-ISG collection, cells maintain their ability to respond to IFN-I. A549s were transfected with modISG10, stimulated with IFNα2b (1000 IU) or left untreated. Cells were then rested for 24hrs and stimulated for 30min with IFNα2b. Levels of pSTAT1, pSTAT2, USP18 and GAPDH were measured by western blot.

**Figure S4.**
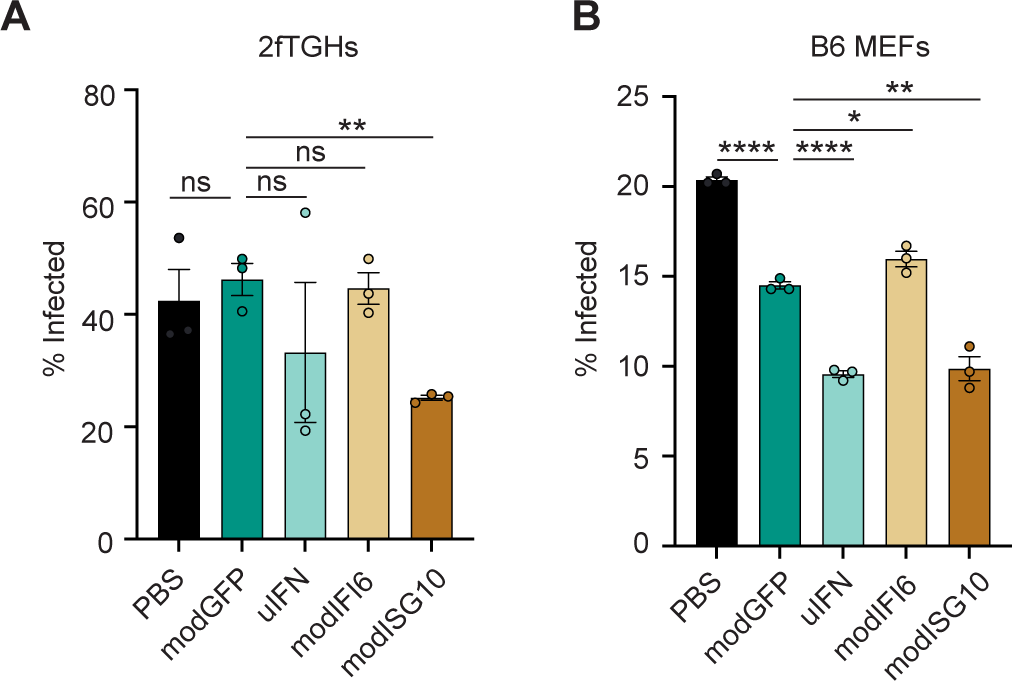
Treatment with synthetically modified collection of 10 ISGs is sufficient to restrict West Nile virus in vitro. B6 MEFs and 2fTGHs were transfected with modGFP, modISG10, modIFI16, stimulated with uIFN (100IU) or left untreated. Cells were then infected with West Nile virus for 24hrs. The percentage of infected (**A**) 2fTGHs or (**B**) B6 MEFs was quantified by staining for flavivirus E-protein using image cytometry.

**Figure S5.**
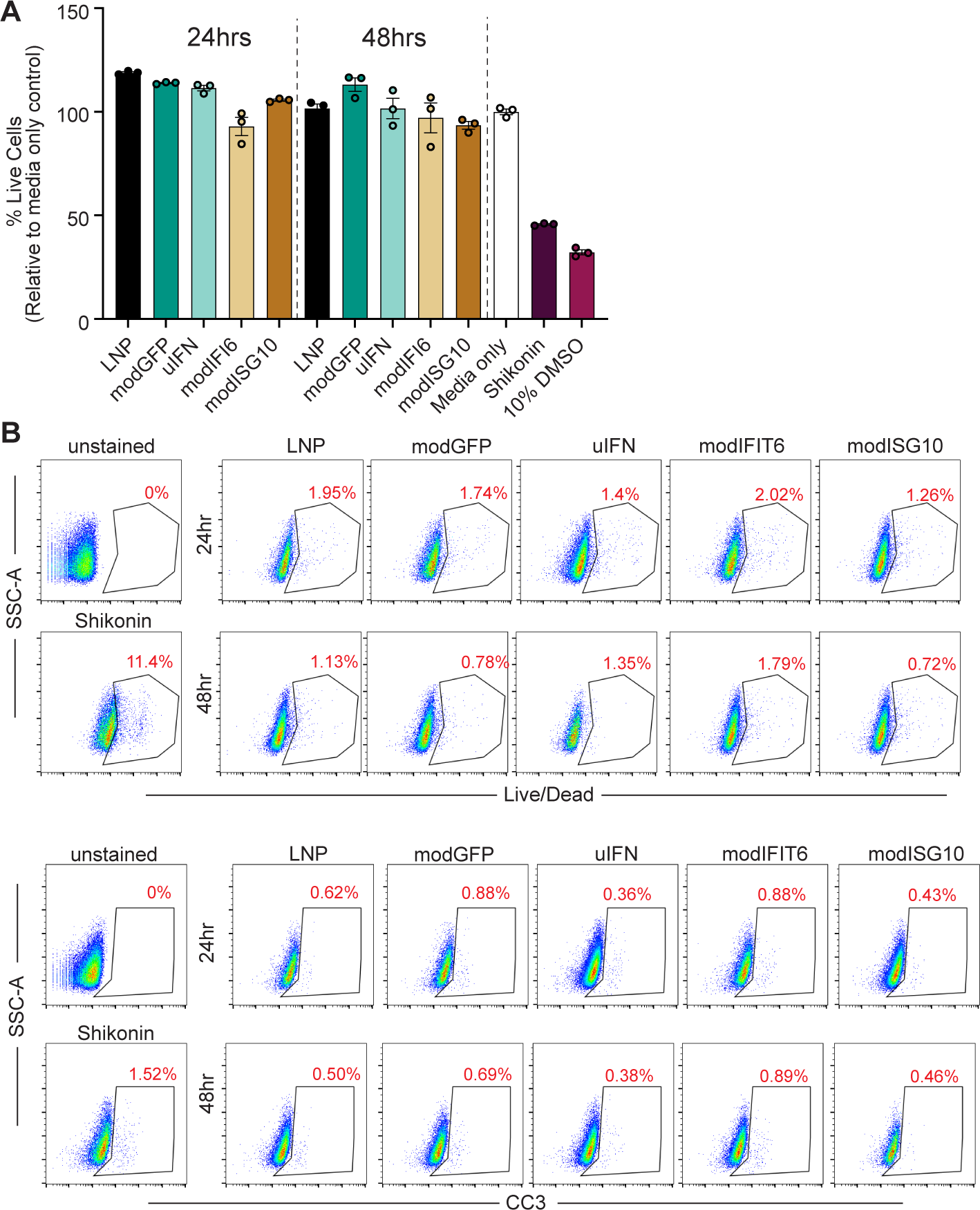
Delivery of packaged modified 10-ISG RNA does not induce cell death *in vitro*. (**A**) Relative viability of hTERT-immortalized fibroblasts following treatment with RNAiMax Lipofectamine (LNP alone), modGFP, modIFI6, modISG10 or uIFN (100IU/ml) was measured with Deep Blue assay. Cells were also treated with either 10μM Shikonin or 10% DMSO to induce cell death. (**B**) The percentage of total dead cells (top) and cleaved caspase positive cells (bottom) was evaluated by flow cytometry.

**Figure S6.**
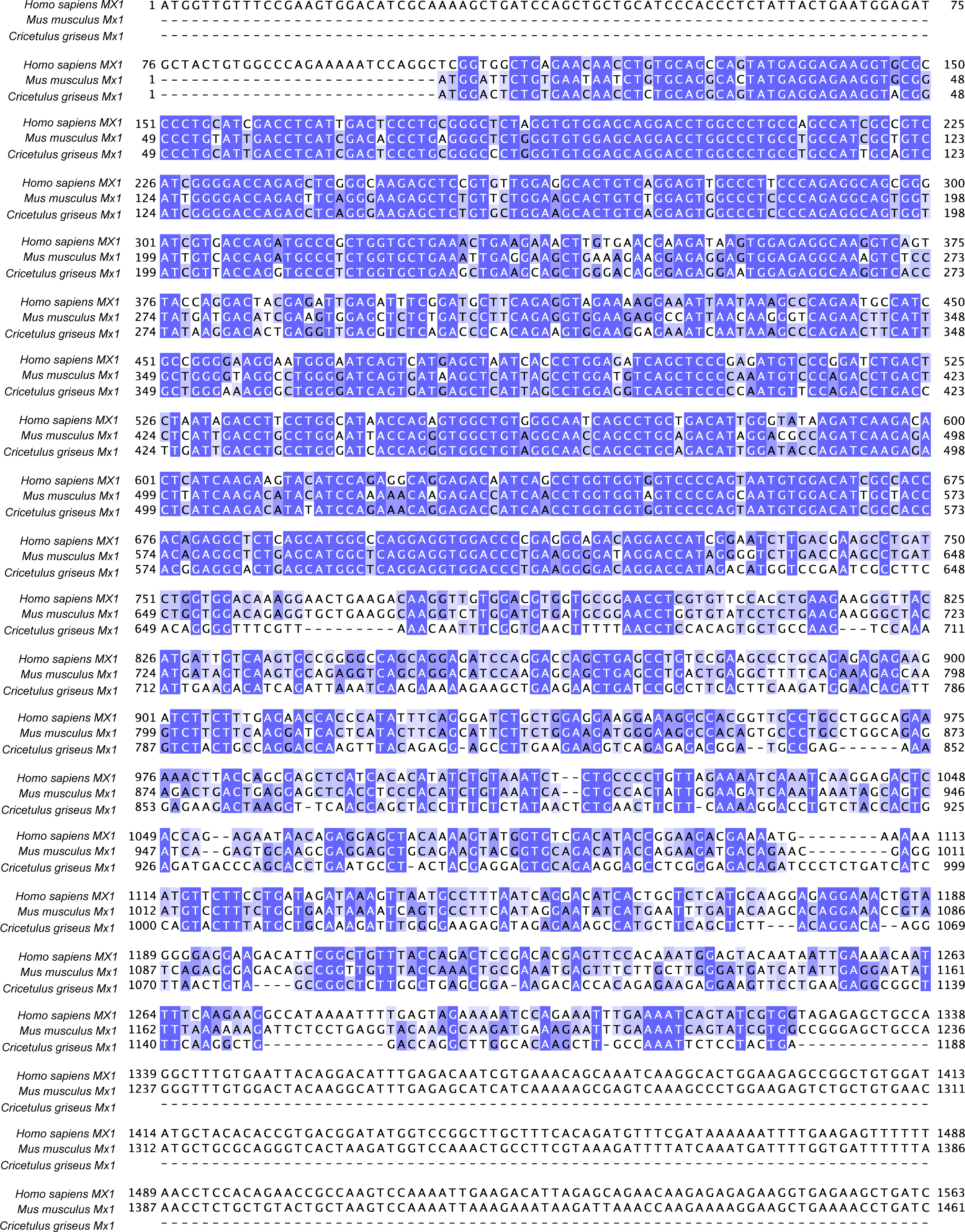
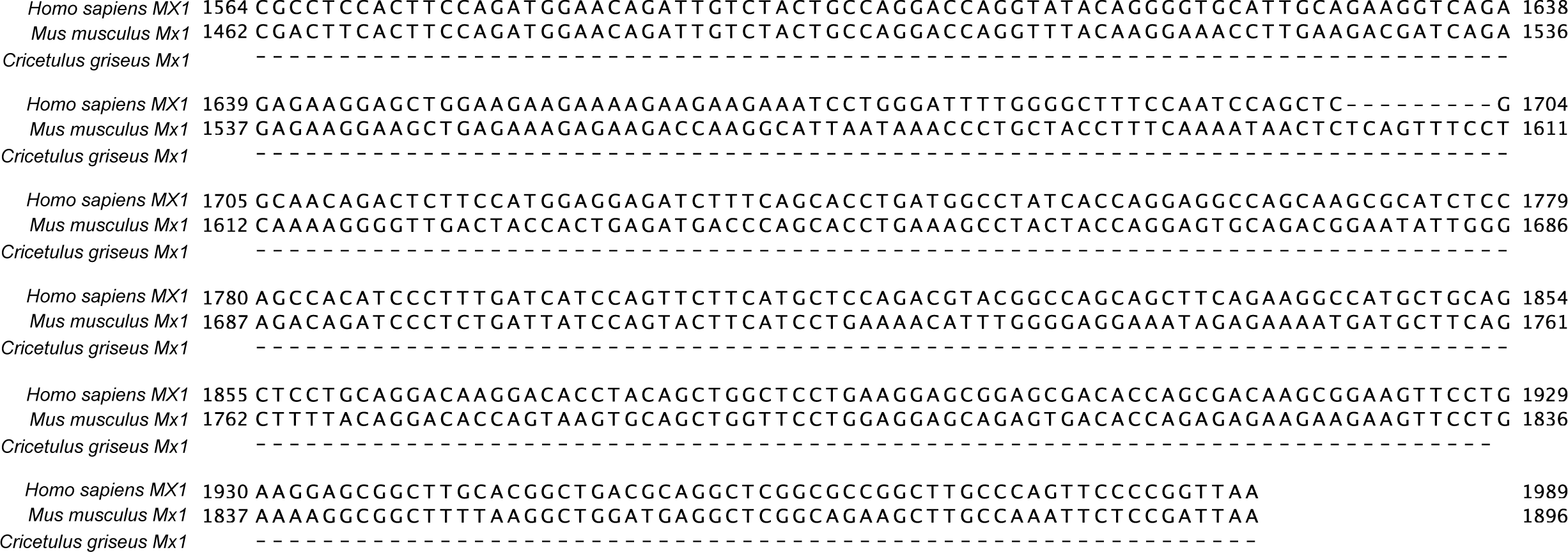
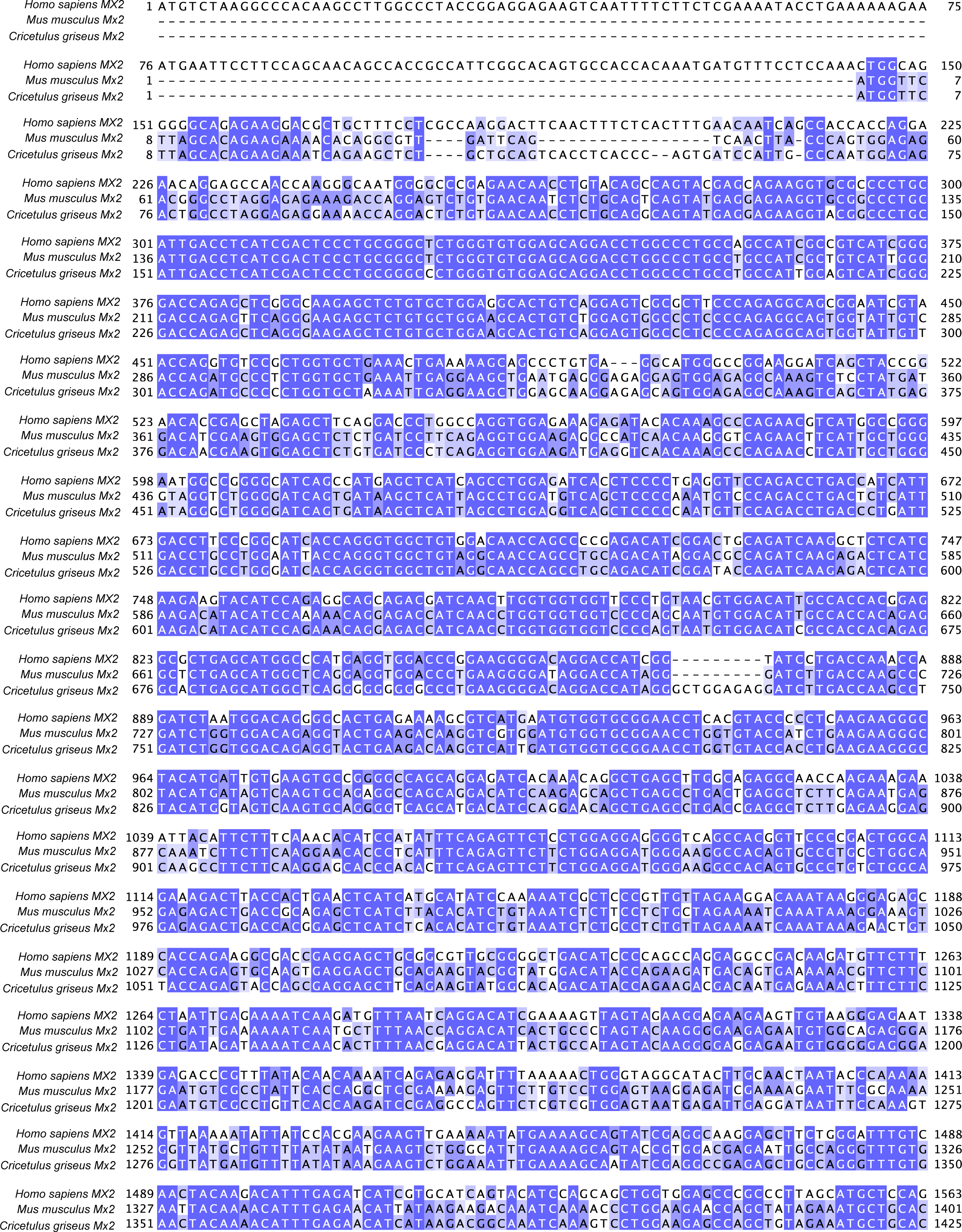
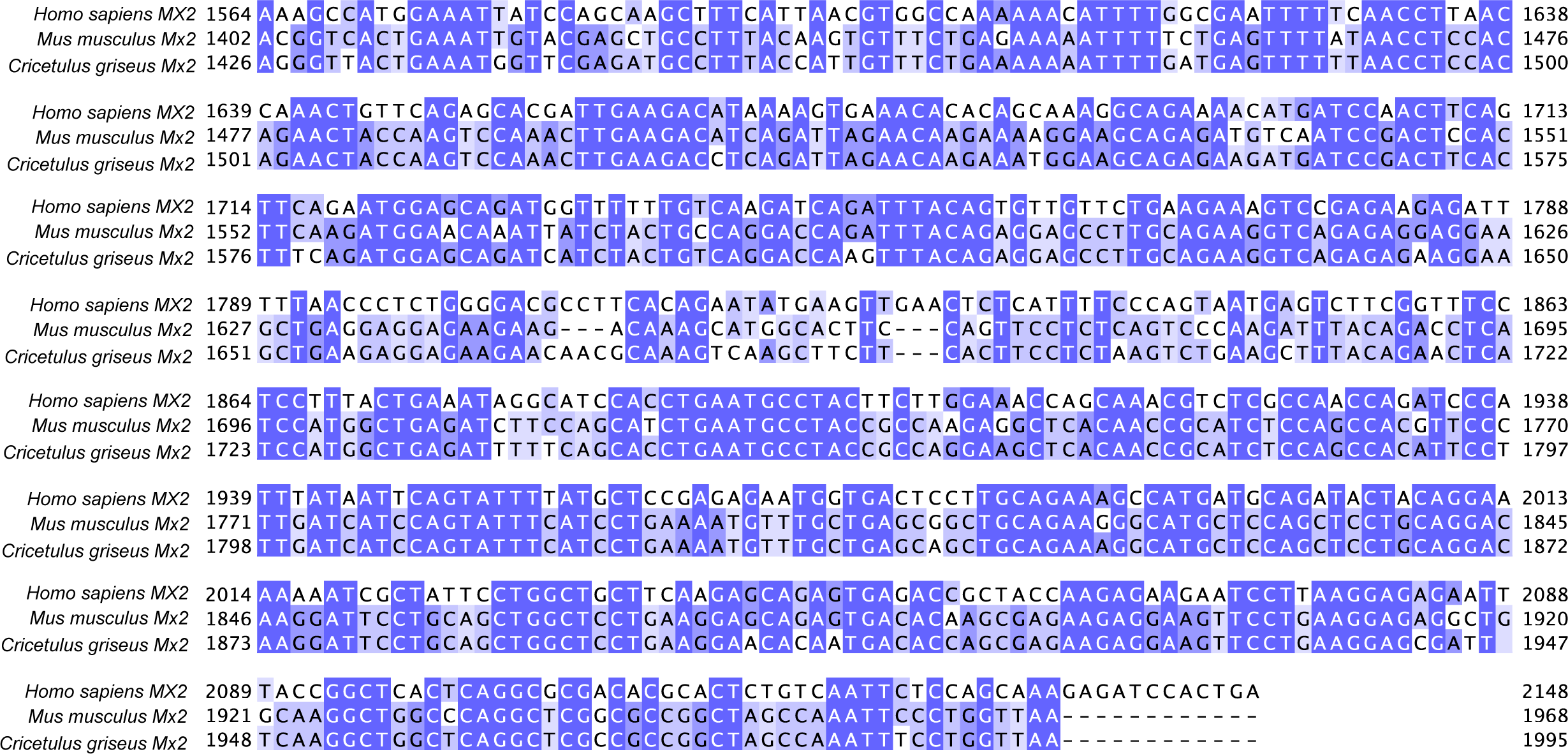
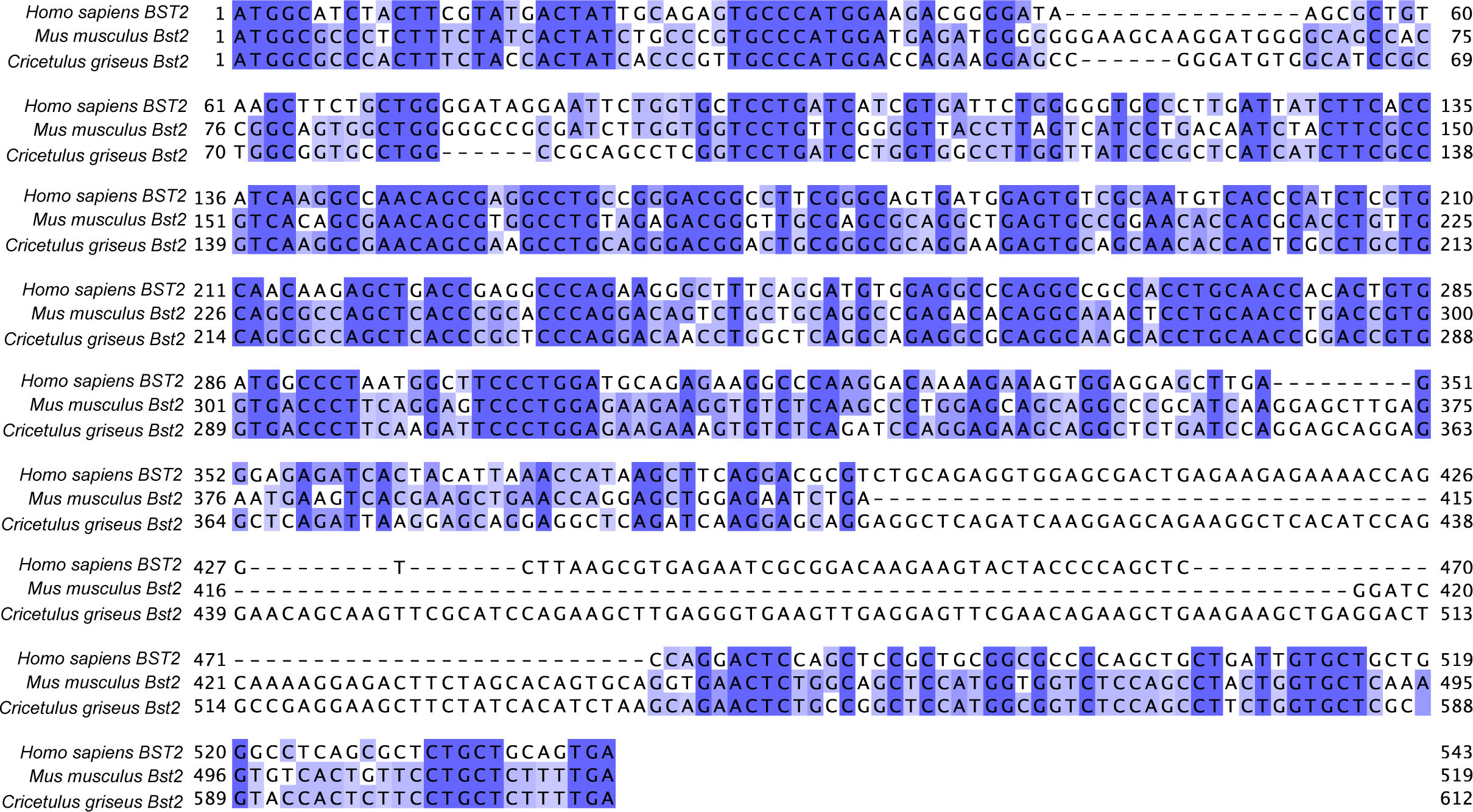
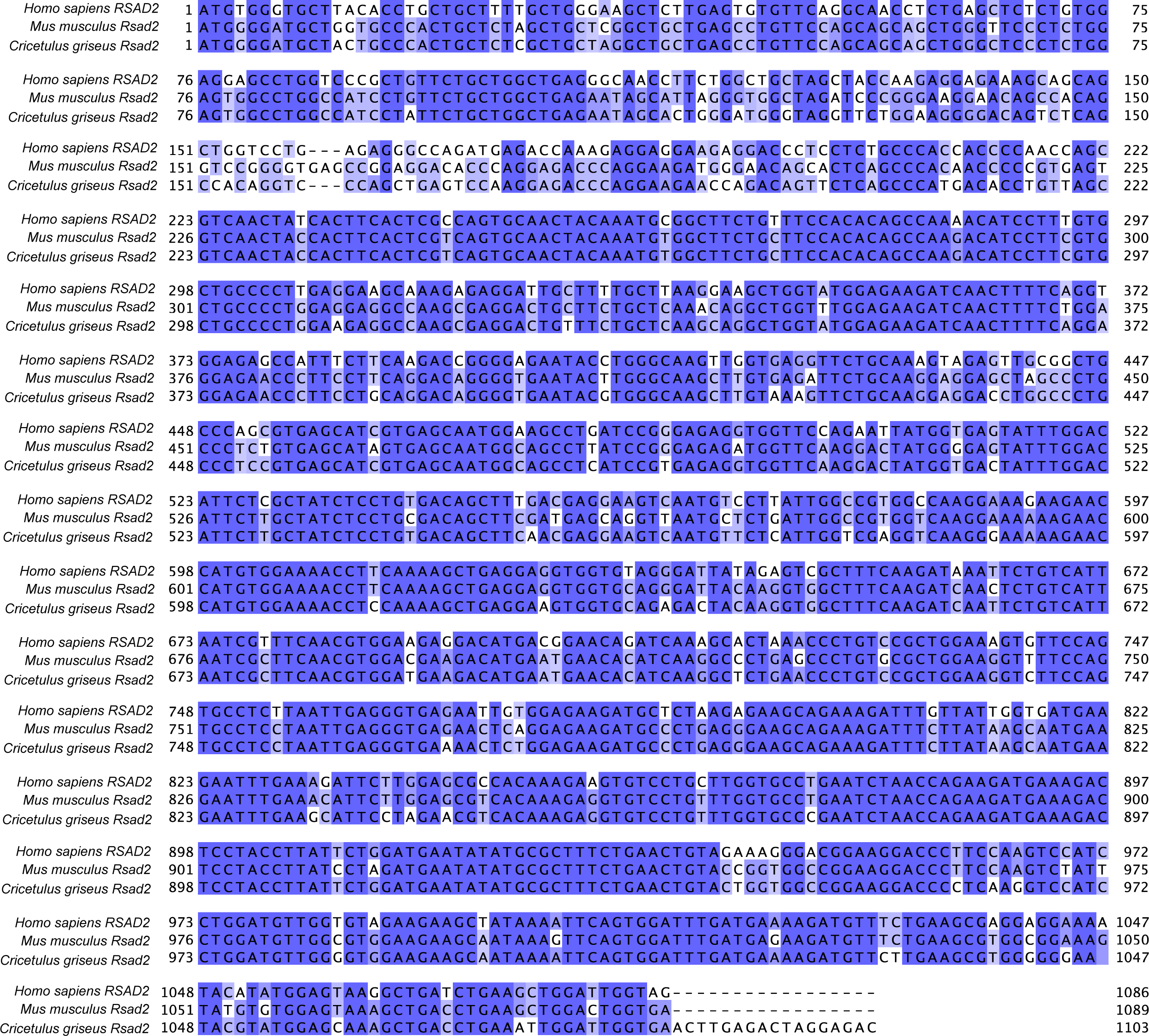
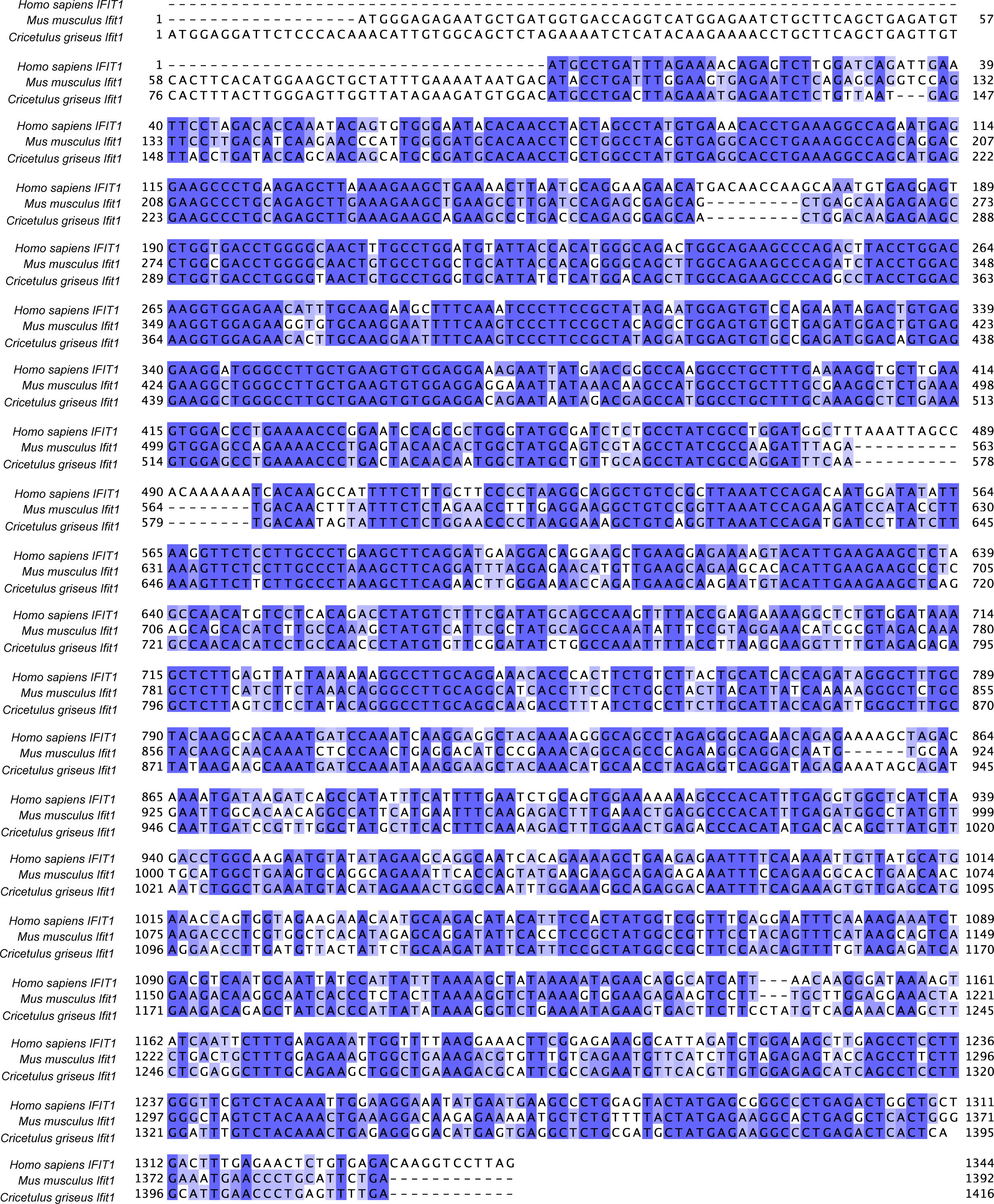
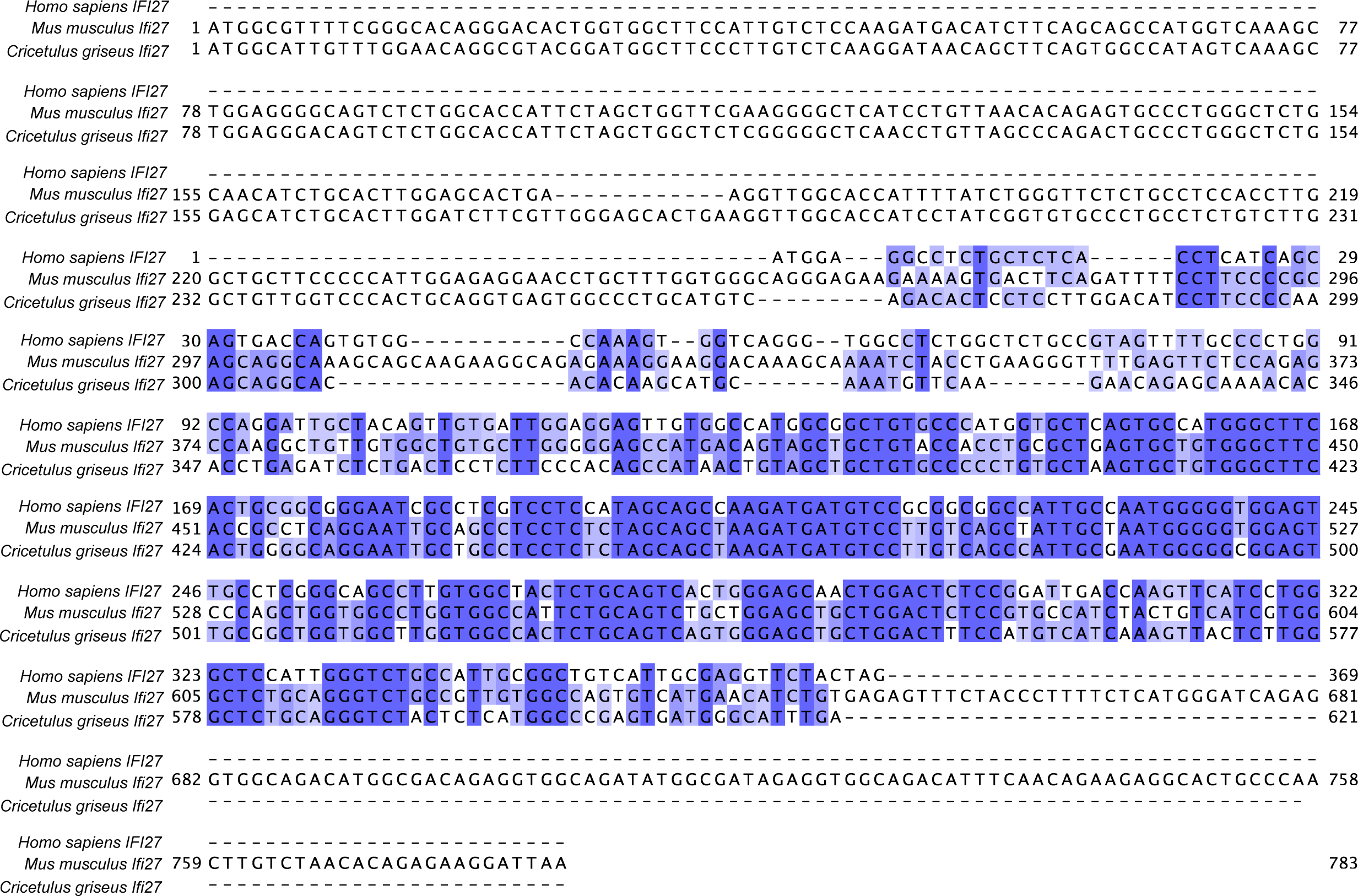
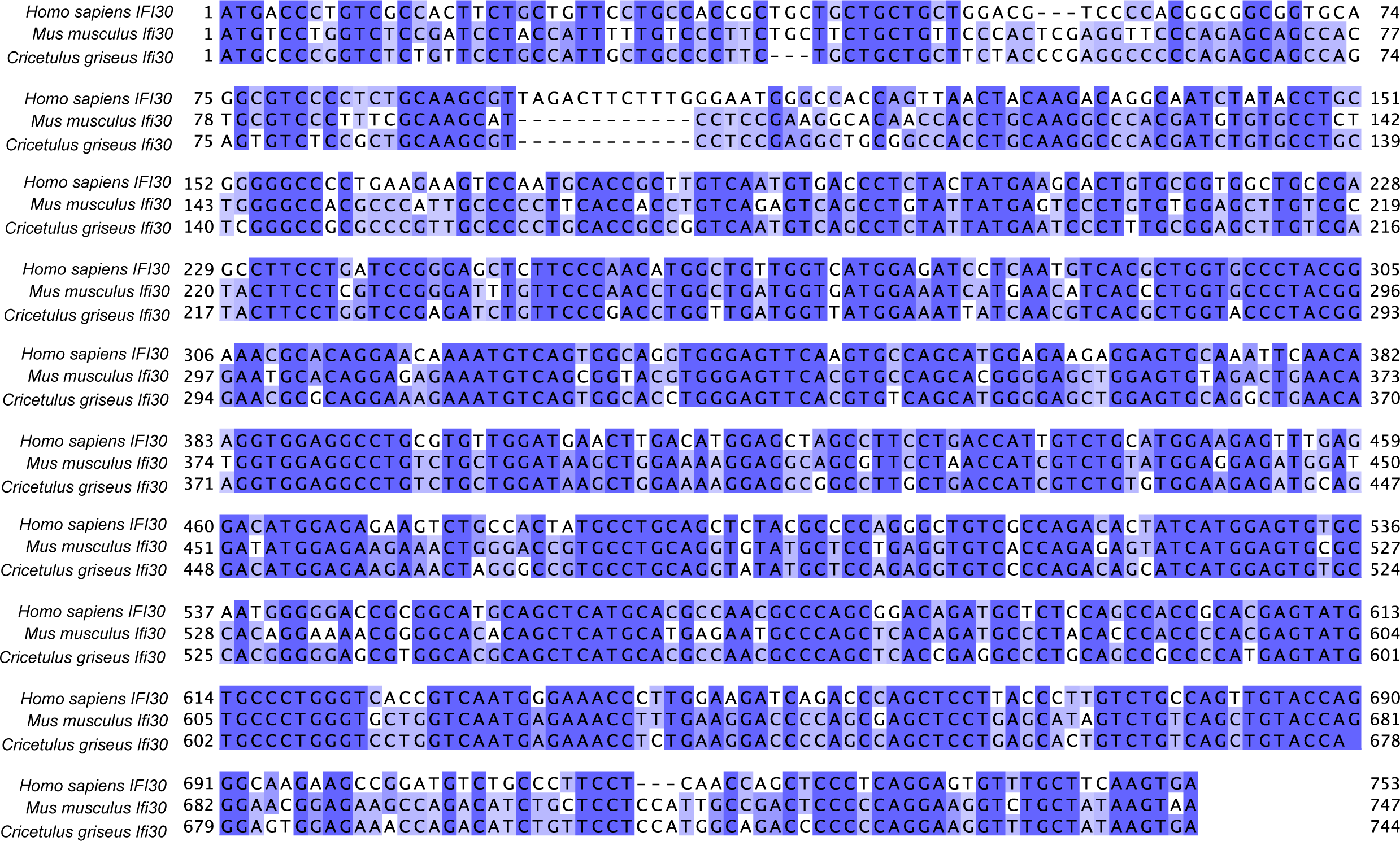
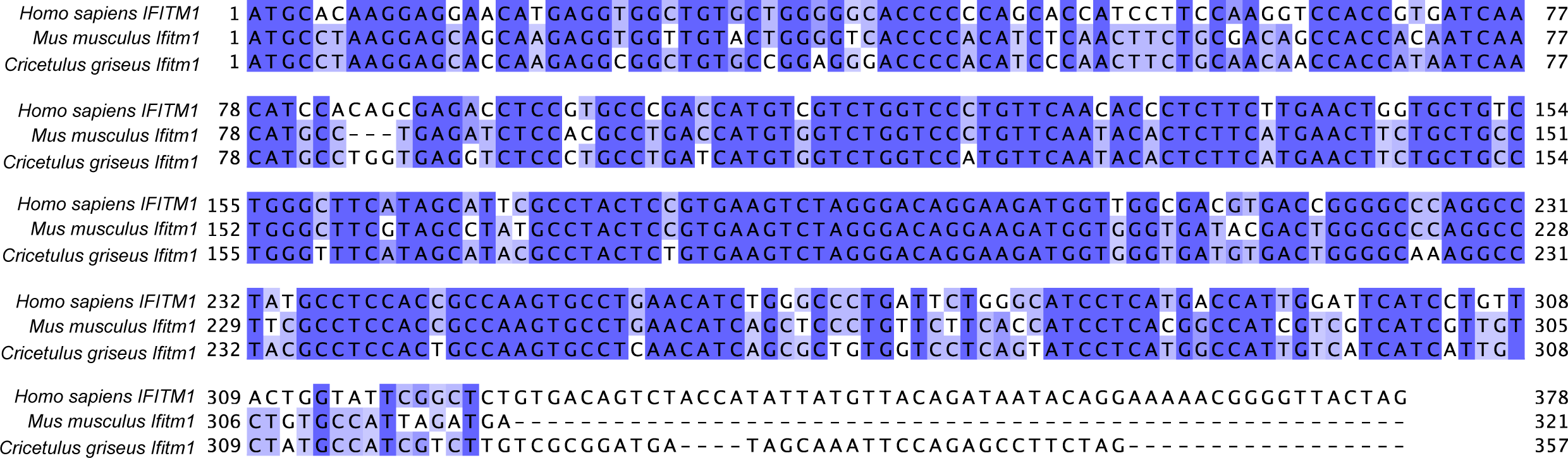
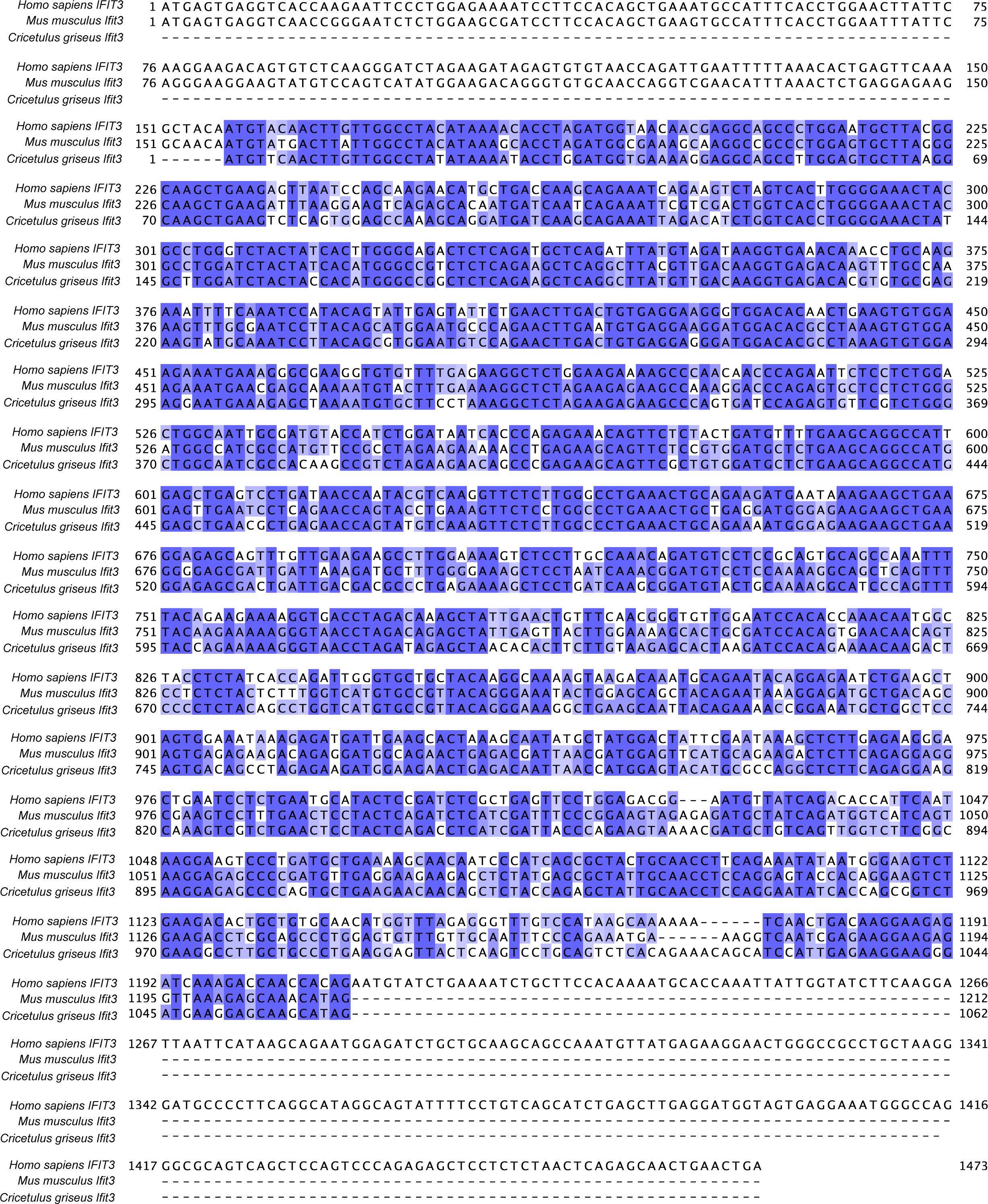
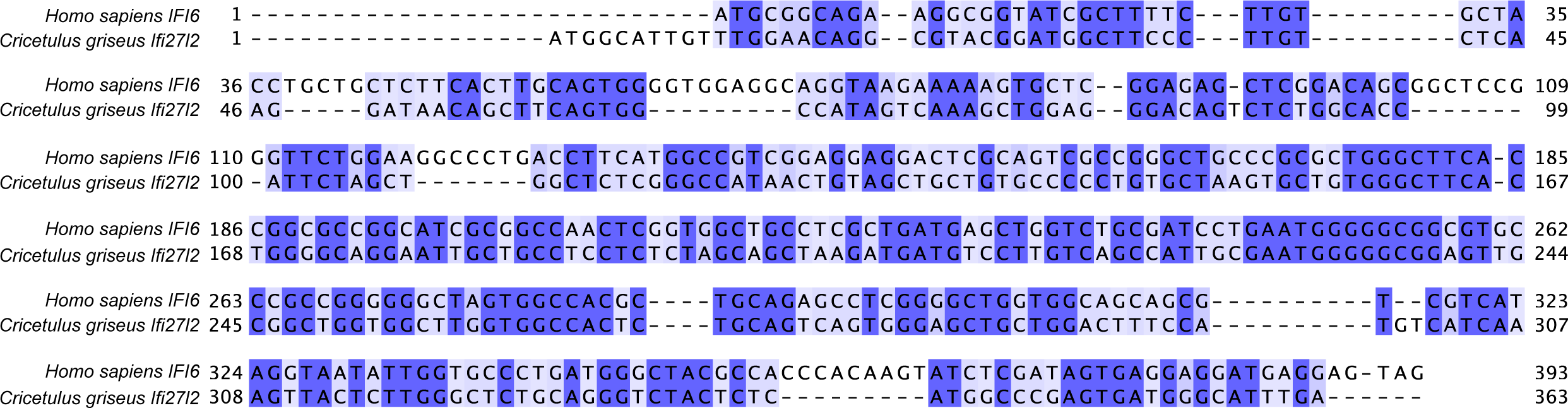
Sequence homology of the nominal collection of 10 ISGs across human, mouse and hamster. Multiple sequence alignment of the open reading frames for the 10 ISGs was generated using Clustal Omega, comparing the orthologues from human (*Homo sapien*), mouse (*Mus musculus*) and Chinese hamster (*Cricetulus griseus*).

**Figure S7.**
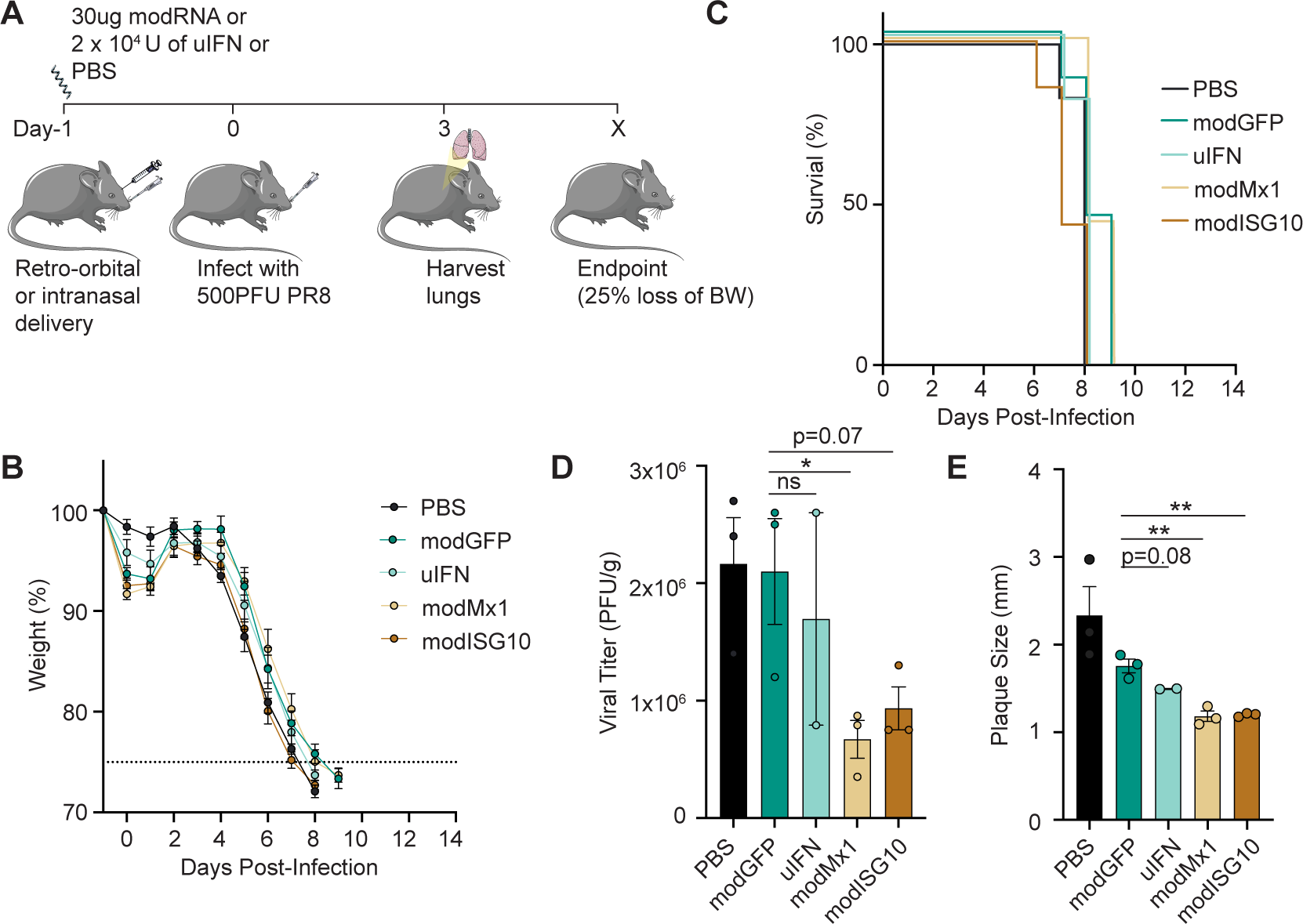
Retro-orbital delivery of synthetically modified RNA of mouse Mx1 or the collection of 10 ISGs significantly reduces lung viral titers in mice challenged with lethal dose of Influenza A virus. (**A**) Schematic showing retro-orbital delivery of 30μg of modGFP/modMx1/modISG10 with Dlin-MC3 DMA:DSPC:Cholestrol:DMG-PEG in C57BL/6 mice on day -1. Alternatively, mice were given 2 x 10^4^ PFU of uIFN intranasally. Control mice were given PBS either retro-orbitally or intranasally. On day 0, all mice were intranasally challenged with 500PFU of PR8 strain of H1N1 influenza virus. (**B**) Mice were monitored for body weight loss (**C**) Mice were considered to have reached the end point when they lost 25% of their initial body weight. The broken line in (**B**) indicates 75% of initial body weight. (**D-E**) At day 3 post-infection, lungs were harvested from n=2-3 mice per group to perform a plaque assay (*p<0.05, **p<0.01, unpaired Student’s t-test).

## Methods

### Cell culture

Icahn School of Medicine IRB board has approved the use of human subject cell lines. Dermal fibroblasts from *ISG15*^-/-^ patients (*n*=3) and WT controls (*n*=3) were immortalized by stable transduction with human telomerase reverse transcriptase. *ISG15*^-/-^ fibroblasts harbor either a nonsense or frame-shift mutation at position c.379 or c.336_337, respectively. As mutations are in the last exon of *ISG15*, these transcripts are not subject to RNA decay but result in total loss of protein expression (Bogunovic et al., 2012; Zhang et al., 2015). HEK293T (CRL-11268; ATCC), Vero-E6 (CRL-1586; ATCC), Vero (CCL81; ATCC), U4C JAK1^-/-^ (gift from Sandra Pelligrini), 2fTGH (JAK1^+/+^ U4C parental; a gift from Sandra Pelligrini), A549 (CCL-185; ATCC) and human fibroblasts were cultured in normal growth medium consisting of DMEM supplemented with 10% fetal calf serum, 1% L-Glutamine and 1% penicillin/streptomycin. All cells were cultured at 37 °C and 10% CO_2_. All cells were tested for mycoplasma contamination using the PlasmoTest kit according to the manufacturer’s instructions (Invivogen).

### Animals

Six-to eight-week-old female C57BL/6 mice (stock: 000664) were purchased from Jackson Laboratories. Four-to six-week-old female Syrian golden hamsters (Crl:LVG(SYR); strain code: 049) were purchased from Charles River Laboratories. The animals were maintained under a strict 12-hour light cycle and fed a standard chow diet *ad libitum* in a specific pathogen-free barrier facility at Icahn School of Medicine at Mount Sinai (ISMMS). All experiments involving animals were performed in accordance with the regulatory guidelines and standards set by the Institutional Animal Care and Use Committee of ISMMS.

### Lentivirus generation and relevant ISG expressing cell lines

Single ISG ORFs encoding human *MX1, MX2, IFIT1, IFIT3, IFITM1, IFI6, IFI27, IFI30, BST2,* and *RSAD2* (Table S1) were independently cloned from an IFN-a2b stimulated control fibroblast line and inserted into the pTRIP vector backbone by gateway ligation, described previously (Schoggins et al., 2011). To generate lentiviruses, 2.5 μg of the desired transfer vector, 2 μg psPAX2 and 0.8 μg of pMD2.G, 14 μl Lipofectamine 3000 (Fisher scientific #L3000001), and 20μl of Lipofectamine P3000 reagent, was prepared in 250μl of OptiMEM (Gibco #11058021). This transfection mix was gently added by dripping on to 1.5×10^6^ HEK293T cell monolayer in 2 ml of 10% FBS DMEM for 16 hours. Transfection media was removed and replaced with 2 ml of 10% FBS DMEM. 48 hours post-transfection, lentivirus supernatants were collected and centrifuged at 1000 x *g* (RCF) for 5 minutes to pellet cell debris and stored at -80 °C. Collected lentiviruses were each transduced into U4C JAK1^-/-^ cells with single ISG lentiviruses for 24 hours in the presence of polybrene (8 μg/ml, Signa #H9268), before puromycin antibiotic selection (0.4 μg/ml). Cells remained under selection until all cells in the non-transduced cells were floating/dead. Successful transductants were confirmed for RFP expression and stored at -80°C.

### Synthetic modified RNA synthesis

Production of in vitro transcription (IVT) template constructs and subsequent RNA synthesis have been described previously (Warren et al., 2010; Zangi et al., 2013). All oligonucleotide reagents were synthesized by Integrated DNA Technologies (Coralville). ORFs were amplified by PCR from plasmids encoding emerald GFP, firefly luciferase, and human ISGs (Table S1) or their hamster (Table S2) or mouse (Table S3) orthologs. Due to the Syrian golden hamster genome reference being incompletely annotated, hamster orthologs were selected by querying the Chinese Hamster and/or the Chinese hamster ovary cell line genome reference using the Ensembl API (v100). For *IFI6*, the closest homolog sequence (*Ifi27l2*) was chosen (Table S2). According to the manufacturer’s instructions, PCR reactions were performed with HiFi Hotstart (KAPA Biosystems). Splint-mediated ligations were carried out with Ampligase Thermostable DNA Ligase (Epicenter Biotechnologies). UTR ligations were conducted in the presence of 200 nM UTR oligos and 100 nM splint oligos. All intermediate PCR and ligation products were purified with QIAquick spin columns (Qiagen) before further processing. Template PCR amplicons were subcloned with the pcDNA 3.3-TOPO TA cloning kit (Invitrogen). Plasmid inserts were excised by restriction digest and recovered with SizeSelect gels (Invitrogen) before being used to template Poly A tail PCRs. RNA was synthesized with the MEGAscript T7 kit (Ambion), with 1.6 μg of purified tail PCR product to template each 40 μl reaction. A custom ribonucleoside blend was used comprising 3′-O-Me-m7G(5′)ppp(5′)G cap analog (New England Biolabs), ATP, and guanosine triphosphate (USB), 5-methylcytidine triphosphate and pseudouridine triphosphate (TriLink Biotechnologies). Final nucleotide concentrations in the reaction mixture were 6 mM for the cap analog, 1.5 mM for guanosine triphosphate, and 7.5 mM for the other nucleotides. RNA was purified with Ambion MEGAclear spin columns and then treated with Antarctic Phosphatase (New England Biolabs) for 30 min at 37 °C to remove residual 5′-phosphates. Treated RNA was repurified, quantified by Nanodrop (Thermo Scientific), and precipitated with 5 M ammonium acetate according to the manufacturer’s instructions. modRNA was resuspended in 10 mM Tris HCl, 1 mM EDTA at 100 ng/μl for in vitro use or 20–30 μg/μl for in vivo use.

### Lipid nanoparticle packaging and transfection

modRNA and RNAiMAX (Invitrogen) transfection agent were each dissolved separately in Opti-MEM (Invitrogen), combined, and then incubated for 15 min at room temperature to generate the transfection mixture. For the 10-ISG collection treatments, modRNA for each ISG were equimolarly pooled before packaging within RNAiMAX lipid reagent. 5 or 0.5 μl of RNAiMAX reagent was used for every microgram of modRNA for in vitro or in vivo transfection in hamsters. In vitro transfection was performed by adding the transfection mixture to cells plated in DMEM with 2% FBS (eBioscience, San Diego, CA). For in vivo transfection of hamster lungs, the transfection mixture was administered intranasally via a liquid drip into the nasal cavity or intratracheally via injection. Alternatively, modRNAs administered in mice were packaged in Dlin-MC3-DMA formulated with 1,2-distearoyl-*sn*-glycero-3-phosphocholine (DSPC), cholesterol, DMG-PEG and DOTAP as previously described (Cheng et al, 2020). Dlin-MC3-DMA: DSPC: cholesterol: DMG-PEG was prepared with a molar ratio of 50:10:38.5:1.5 with 50% DOTAP.

### IFN-I cell stimulation and exposure experiments

Control or ISG15^-/-^ fibroblasts were stimulated with 0 or 1,000 IU ml^−1^ IFN-α2b (Merck IntronA) in normal growth medium for 12 h. The IFN-α2b was then eliminated by thorough washing (3X) with PBS, and the cells were allowed to rest for 36 h in normal medium before infection or being passed off to other downstream assays.

### RT-qPCR

Cell lines were lysed, and RNA was isolated with Rneasy spin columns (QIAGEN 74104). Reverse transcription was performed with the High-Capacity RT Kit (Applied Biosystems 4368814). The resulting cDNA was then subjected to qPCR with the TaqMan Master Mix II with UNG (Thermo Fisher 4440038), on a Roche LightCycler 480 or Applied Biosystems QuantStudio 6, with the following primers/probes: 18S(4318839), MX1 (Hs00895608), RSAD2 (Hs00369813), SIGLEC1 (Hs00988063), IFIT1 (Hs01911452), BST2 (Hs00171632), USP18(Hs00276441), ISG15 (Hs01921425), IFI44L (Hs00915292), GAPDH (Hs01922876). The relative expression of each transcript was normalized relative to 18S or GAPDH by the ΔΔCt method.

### Mass spectrometry

Cells were washed in ice-cold PBS and collected in PBS. Cell pellets were lysed in 100μL of 8M Urea, 50mM ABC, 150mM NaCl, and Halt Protease Inhibitors (78429) in Fisher Gold HPLC water (W64). Protein concentration was measured with the Thermo microBCA kit (23235). 20μg of lysate was removed from each condition. Protein was then reduced with tris (2-carboxyethyl) phosphine (TCEP) to a final concentration of 2mM (PG82089). Following reduction, proteins were alkylated with iodoacetamide (IAA) to a final concentration of 10mM (i6125-5g). Excess IAA was quenched with dithiothreitol (DTT) to a final concentration of 10mM (D5545-1G). Four volumes of 50mM ammonium bicarbonate were added to dilute the urea to less than 2M. Proteins were digested with a 1:100 trypsin to protein ratio. Digest took place over 16 hours with end-over-end rotation at room temperature. Samples were acidified to 0.03% TFA to stop the digestion. Samples were then desalted according to manufacturer’s protocol using Nest Group C18 spin columns (HUM S18R) and dried by vacuum centrifugation. All samples were analyzed on an Orbitrap Eclipse mass spectrometry system (Thermo Fisher Scientific) equipped with an Easy nLC 1200 ultra-high pressure liquid chromatography system (Thermo Fisher Scientific) interfaced via a Nanospray Flex nanoelectrospray source. For all analyses, 1mL of sample containing 0.5ug of digested peptides was injected onto a C18 reverse phase column (30 cm x 75 mm ID) packed with ReprosilPur 1.9 mm particles. The mobile phase A consisted of 0.1% formic acid (FA), and mobile phase B consisted of 0.1% FA/80% acetonitrile (ACN). Peptides were separated by an organic gradient from 5% to 35% mobile phase B over 120 minutes, followed by an increase to 100% B over 10 minutes at a flow rate of 300 nL/min. The analytical columns were equilibrated with 3 mL of mobile phase A. For the spectral library, samples from each set of biological replicates were pooled and analyzed with a Data Dependent Analysis (DDA) Method. The DDA was performed by acquiring a full scan over an m/z range of 375-1025 in the Orbitrap at 120,000 resolving power (@200 m/z) with a normalized AGC target of 100%, an RF lens setting of 30%, and an instrument-controlled ion injection time. Dynamic exclusion was set to 30 seconds with a 10-ppm exclusion width setting. Peptides with charge states 2-6 were selected for MS/MS interrogation using higher energy collisional dissociation (HCD) with a normalized HCD collision energy of 28%, with three seconds of MS/MS scans per cycle. Data-independent analysis (DIA) was performed on all individual samples. An MS scan at 60,000 resolving power over a scan range of 390-1010 m/z, an instrument controlled AGC target, an RF lens setting of 30%, and an instrument-controlled maximum injection time, followed by DIA scans using 8 m/z isolation windows over 400-1000 m/z at a normalized HCD collision energy of 28%. Peptides and proteins were first identified with Spectronaut as previously described (Bruderer R., et al, 2017). False discovery rates were estimated using a decoy database strategy. All data were filtered to achieve a false discovery rate of 0.01 for peptide-spectrum matches, peptide identifications, and protein identifications. Search parameters included a fixed modification for carbamidomethyl cysteine and variable modifications for N-terminal protein acetylation and methionine oxidation. All other search parameters were set to the defaults for the respective algorithms. Analysis of protein expression was conducted utilizing the MSstats statistical package in R. Output data from Spectronaut were annotated based on the human reference (20191010.SwissProt.Hsapiens.fasta). Technical and biological replicates were integrated to estimate log_2_ fold changes, p values, and adjusted p values (Choi, M. 2014). All data were normalized by equalizing median intensities, the summary method used was Tukey’s median polish, and the maximum quantile for deciding censored missing values was 0.999. Significantly dysregulated proteins were defined as those which had a fold change value >2 or <-2, with a p-value of <.0.05.

### SARS-CoV-2 virus propagation and virus titration

A working stock of SARS-COV-2 (isolate USA-WA1/2020, BEI resource NR-52281) was prepared by low multiplicity of infection of Vero-E6 cells with SARS-CoV-2 isolate (passage 2) as previously described (Amanat et al., 2020). Infected cultures were maintained in reduced-serum DMEM (2% FBS) for 4 days before preparation of storage of virus aliquots at -80°C. Viral titers, reported as plaque-forming units Propagation of SARS-CoV-2 propagations and experiments were performed in the BSL-3 Conventional Biocontainment Facility, a Dean’s CoRE facility, in compliance with the approval by the ISMMS Institutional Biosafety Committee (IBC).

### In vitro viral infections

Cells were infected with VSV-GFP, IAV PR8-GFP, ZIKV, or SARS-CoV-2 virus at indicated MOIs for the indicated time (detailed in Figures 2-3). Following a 1-hour infection incubation, cells were washed to remove the input virus, and the infection was allowed to continue for the indicated amount of time. At the end of the infection experiment (detailed in Figures 2-3), cells were fixed in 4% PFA for 20 min at 4°C and stained with DAPI. For fluorescent viruses (VSV-GFP and IAV PR8-GFP), cells were imaged for GFP and DAPI fluorescence signals. For non-fluorescent viruses (ZIKV and SARS-CoV-2), cells were permeabilized and subsequently stained with an antibody targeting the viral envelope protein (Clone 4G2) for ZIKV and the virus nucleoprotein (gift from Thomas Moran) for SARS-CoV-2. GFP signal from infected cells was acquired on a Celigo Imaging Cytometer (Nexcelom) and quantified in CellProfiler (McQuin, et al., 2018). CellProfiler was used in all cases for quantification (McQuin, et al., 2018). DAPI was segmented (global, Minimum Cross-Entropy) to measure percent infection to provide a total cell count. GFP was segmented (global, Minimum Cross-Entropy) to define the entire infected area. Nuclei that overlapped with the infected area were considered infected.

### In vivo viral infections

Syrian golden hamsters were anesthetized with ketamine/xylazine prior to experimental manipulation and euthanized by intracardiac administration of SleepAway euthanasia solution. Experiments with SARS-CoV-2 infected hamsters were performed in an ABSL-3 biocontainment facility at ISMMS. Hamsters were intranasally inoculated with 1×10^5^ PFU (titered on Vero-E6; diluted in 100μl PBS) of SARS-CoV-2 virus. Similarly, mice were anesthetized with ketamine/xylazine (95mg/kg ketamine and 10mg/kg xylazine) and intranasally inoculated with 30μl PBS containing 500PFU (titrated on MDCK cells) of IAV strain A/PR/8/34 (HINI) (PR8) virus.

### TCID_50_/ml viral titer quantification

TCID_50_ viral titrations were conducted as previously described in (Amanat et al., 2020; Reed et al., 1938). Briefly, supernatants from infected wells were collected and spun at 1000 x *g* (relative centrifugal forces; RCF) for 10 minutes at 4 °C to remove cell debris and isolate virus. A day before TCID50 assay, indicated adherent cell lines were seeded in a 96 well format to confluency. Collected viral supernatants were serially diluted and incubated on cell monolayers for 1 week. For GFP fluorescent viruses, plates were fixed with 4% PFA, stained with DAPI, and scanned for GFP signal using the Celigo Imaging Cytometer. For non-fluorescent viruses, plates were fixed with 4% PFA, stained with DAPI and relevant viral epitope protein (ZIKV and WNV flavivirus envelope-protein antibody (clone 4G2), SARS-CoV-2 nucleoprotein), and then scanned for antibody signal using the Celigo Imaging Cytometer. TCID50 values were calculated as described in (Reed et al., 1938).

### Plaque assay for quantification of viral titers of tissue specimens

To determine viral titers, 1×10^6^ Vero-E6 were seeded in a 6 well-plate the day before the plaque assay was performed. Briefly, 10-fold serial dilutions were prepared in infection media for SARS-CoV-2 and inoculated onto a confluent Vero-E6 cell monolayer. After 1-hour adsorption, supernatants were removed, and cell monolayers were overlaid with minimum essential media (MEM) containing 2% FBS and purified agar (OXOID) at a final concentration of 0.7%. Cells were then incubated for 3 days at 37°C. Cells were fixed overnight with 4% formaldehyde to inactivate any potential live SARS-CoV-2 virus. The overlay was removed, and cells were washed once with PBS. Plaques were visualized by staining with 1% crystal violet (Sigma # C0775) for 15 minutes. Viral titers were calculated considering the number of cells per well and expressed as plaque-forming units (PFU) normalized by tissue weight. A similar protocol was followed in MDCK cells to quantify the influenza A virus strain A/PR/8/34 (H1N1) (PR8). Briefly, MDCK cells were seeded in a 12-well plate the day before the plaque assay was performed. Ten-fold serial dilutions were performed in infection media for IAV PR8 and inoculated onto a confluent MDCK cell monolayer. After 1 hour adsorption, supernatants were removed, and cell monolayer were overlaid with MEM containing 0.1% NAHCO_3_, 0.05% DEAE-DEXTRAN and purified agar (OXOID) at a final concentration of 0.6%. Cells were then incubated for 2 days at 37°C in 6% CO_2_. Cells were fixed with 2% formaldehyde solution, followed by staining with 1% Crystal violet (Sigma # C0775) and viral titers were expressed as PFU per gram of tissue.

### Cell death assay

Fibroblasts were plated at approximately 80-90% confluency in a 96-well plate and kept overnight in growth media containing 10% FBS. As a positive control for test conditions, 10 μM of Shikonin (Santa Cruz Biotechnology #517-89-5) or 10% DMSO was added to induce cell death. Following a 24 or 48-hour incubation, viability was measured using the Deep Blue Cell Viability Kit (Biolegend 424702). Alternatively, cells were collected and stained with Zombie NIR dye (Biolegend 423105) or with anti-cleaved caspase-3 antibody (Biolegend 559341) for quantification by flow cytometry.

### ELISA

Supernatants were collected from A549 cells, 2fTGH and HEK293T cells and ELISA was performed according to the manufacturer’s protocols for human IFN-β (R&D systems DY814-05PBL Assay Science 41410-2), human IFN-γ (Biolegend 43104) and human IFN-λ4 (AFG Bioscience EK716122VWR 76578-602).

### Immunoblots

Cells were lysed in RIPA buffer (Thermo Fisher 89900) containing protease/phosphatase inhibitor cocktail (Cell Signaling 5872) for 20 minutes. Lysates were centrifuged at 14,000rpm for 10 minutes to remove insoluble complexes. Supernatants were boiled with NuPage sample buffer (Thermo Fisher NP0007) containing 50mM DTT. The samples run by gel electrophoresis and proteins were transferred onto PVDF membranes. Membranes were blocked with 5% milk and incubated overnight with primary antibody. Next membranes were incubated with HRP-conjugated secondary antibodies for one hour. Pierce SuperSignal West Pico or Femto Chemiluminescent substrate (Thermo Fisher Scientific 34580/34096) was added to the membranes and bands were visualized using Amersham Imagequant 600 (Cytiva).

### RNA sequencing and relevant analyses

ISG15^-/-^(n=3) and control (n=3) unstimulated and IFN-I exposed (12-hour stimulation; 1000 IU ml-1 IFN-α2b, 24-hour rest) hTERT-immortalized fibroblasts were lysed. Next, total RNA was isolated using the Rneasy spin columns (QIAGEN 74104) and was sent for high-throughput RNA sequencing to Azenta Life Sciences. FASTQ files were uploaded to BaseSpace (Illumina) and aligned to hg19 with the RNA-Seq Alignment (STAR) module. The resulting sample-gene expression counts matrices were exported and read into the R (v4.0.4) statistical computing platform. Metadata and experimental design were extracted from sample names and used to generate a design matrix for differentially gene expression (DGE) analyses. Before DGE analyses, sample-gene matrices were filtered to exclude low expressed genes (counts < 5 in 3 or more samples). DGE analyses were conducted using DESeq2 (v1.30.1) under a paired-sample responsive design (design: ∼genotype + IFN-I stimulation). Resulting DGE lists were filtered against a log_2_FC of 1.5, and known interferon-stimulated genes (Schoggins et al., 2011) were subset from this list. A variance stabilizing normalization function was applied to the sample-gene counts matrix, and differentially responsive ISGs were visualized using the ComplexHeatmap package (v2.11.1) and ggplot2 (v3.3.5). Differentially responsive ISGs between conditions were subject to gene ontology analysis (GO) using the goana function inside the edgeR package (v3.32.1). Briefly, gene symbols were converted into the Entrez IDs by mapping to the human genome annotation database (org.Hs.eg.db; v3.12.1) and fed into the goana function. To mitigate p-value exaggeration and control for cell background specificity, the gene universe was explicitly set only to include genes detected and used for DGE. Resulting GO terms were filtered against a p-value < 1×10^-12^ and subset to only include biological process terms. Passing GO terms were visualized using ggplot2 (v3.3.5).

### Quantification and statistical analysis

All statistical analyses were performed using GraphPad Prism (GraphPad Software Inc) or R. Unless otherwise indicated, error bars represent the standard deviation (SD) from the mean of at least 3 biological replicates. Statistical differences between experimental groups were determined using various statistical tests, including the one-sided paired ratio Student’s t-test, Kaplan-Meier test and one-way ANOVA. A p values of less than 0.05 was considered significant. For differentially gene expression analyses, an adjusted p-value was calculated to account for multiple testing using the Benjamini-Hochberg method.

## Supplementary Information

**Table S1.**
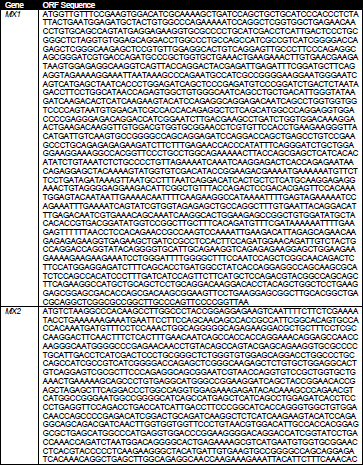
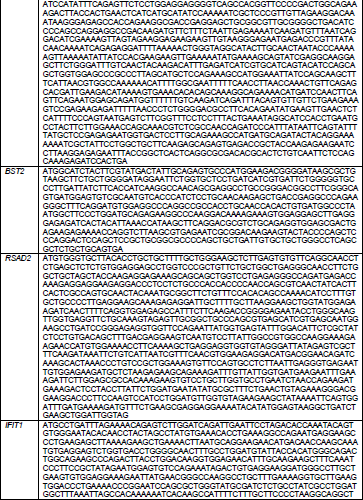
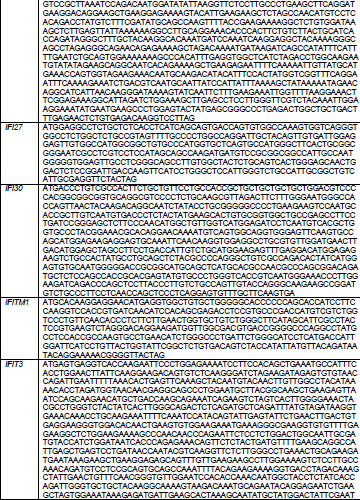
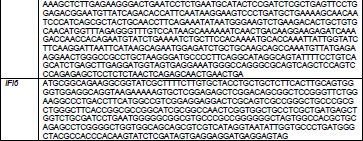
Human (*Homo sapiens*) interferon-stimulated gene sequences.

**Table S2.**
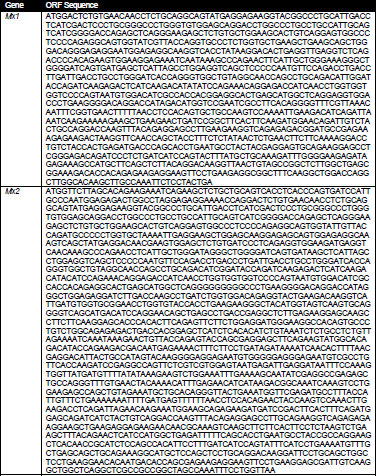
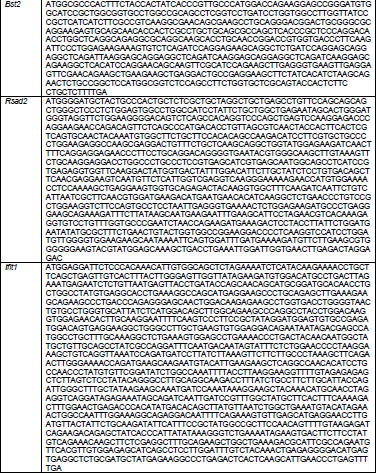
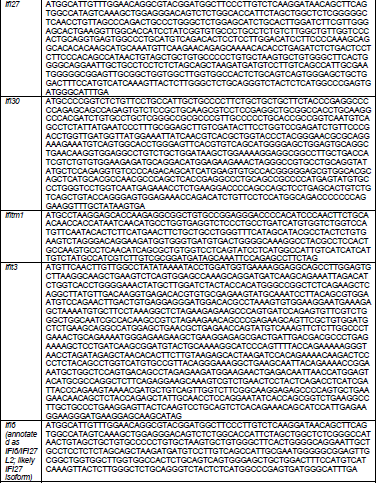
Chinese hamster (*Cricetulus griseus*) interferon-stimulated gene orthologues.

**Table S3.**
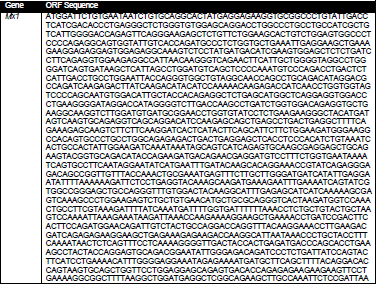
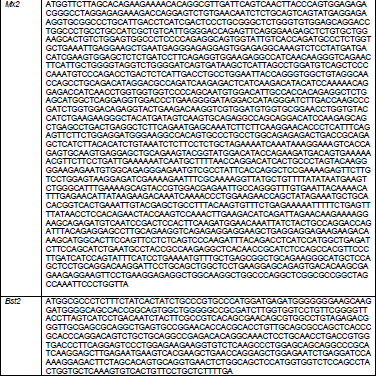
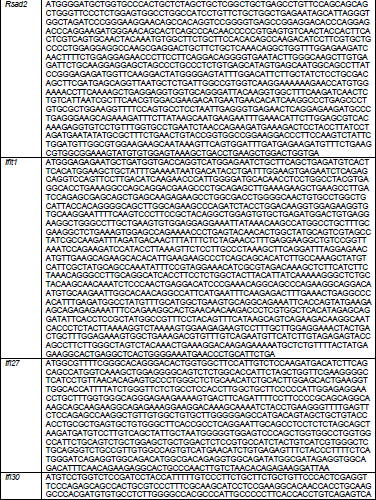
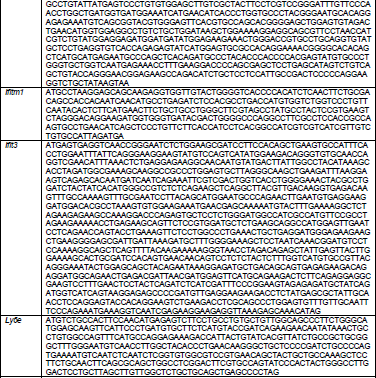
Mouse (*Mus musculus*) interferon-stimulated gene orthologues.

